# Whole-body PET imaging of SIV using anti-Env probes fails to reveal regions of specific uptake in rhesus macaques

**DOI:** 10.1101/2024.07.05.601545

**Authors:** Sharat Srinivasula, Insook Kim, Hyukjin Jang, Paula Degrange, Heather Brown, Viviana Dalton, Yunden Badralmaa, Ven Natarajan, Brad Long, Jorge A. Carrasquillo, Michele Di Mascio

**Author notes:** **Corresponding author:** Dr. Michele Di Mascio.

## Abstract

Following the initial reports demonstrating the feasibility of immunoPET imaging of SIV using anti-Env monoclonal antibodies in non-human primates, replication efforts of the imaging system in HIV-infected individuals have yielded conflicting results. Herein, we used ^89^Zr-7D3 and ^89^Zr- ITS103.01LS-F(ab’)_2_, two anti-gp120 antibodies for immunoPET imaging of SIV in n=20 rhesus macaques. Despite their demonstrated nanomolar affinity and strong binding specificity to SIV gp120 cell lines, we observed no discernible differences in their binding in primary cells, tissue sections of secondary lymphoid organs, *in-vivo* probe uptake between SIVmac-infected and uninfected macaques, or *ex-vivo* validation necropsies. While the probes remained stable *in-vivo*, only ^89^Zr-ITS103.01LS-F(ab’)_2_ in chronic plasma retained its binding specificity to SIV gp120, with ^89^Zr-7D3 experiencing a >97% reduction in binding to gp120 due to competition from endogenous antibodies at the 7D3 binding site. The overall absence of specific uptake suggests inadequate binding potential (ligand affinity x target molarity) for these probes to effectively image SIV or HIV *in-vivo*, warranting further investigation into the lack of reproducibility observed with earlier non-human primate SIV imaging and conflicting human studies.

## Introduction

Global efforts are currently advancing to develop immuno-positron emission tomography (immunoPET) HIV-1 imaging systems that combine the targeting specificity of monoclonal antibody (mAb) and the superior sensitivity and spatial resolution of PET to noninvasively monitor tissue sites of viral replication ^1–7^. Besides their potentially crucial prognostic value of managing people living with HIV (PLWH), these endeavors could also greatly assist in developing strategies for HIV cures and vaccines ^8^.

So far, limited pre-clinical immunoPET studies of the nonhuman primate (NHP) model of HIV-1 pathogenesis reported feasibility in achieving this goal with two different viral envelope (Env) specific mAbs. In a simian immunodeficiency virus (SIV) infected rhesus macaque (RM) model undergoing immunoPET imaging with ^64^Cu radiolabeled PEG-modified intact murine 7D3 mAb (targeting the CCR5 binding site of gp120 envelope glycoprotein ^9^), differences in uptake were reported in various lymphoid tissues and the gastrointestinal tract not only between chronically infected viremic and uninfected but also between antiretroviral therapy (ART) treated aviremic and uninfected adult RMs, thus asserting a remarkably sensitive non-invasive imaging system ^1,2^. In a similar NHP model using ^64^Cu radiolabeled F(ab’)_2_ fragment of primatized 7D3 mAb, viral reactivation in the gut and select lymphoid tissues of long-term ART-treated, aviremic, SIV-infected RMs was reported immediately following administration of a latency reversal agent (LRA), suggesting a sensitive imaging system capable of detecting gp120 in tissues when viral levels are ∼3-4 Log_10_ lower than in chronic viremic animals ^3,4^. In another NHP model of simian/human immunodeficiency virus (SHIV)-infected RMs, ^68^Ga radiolabeled Fab fragment of PGT145 mAb (targeting the variable loop 2 (V2) apex of gp120 ^10,11^) detected viral reactivation in the gastrointestinal tract and some lymphoid tissues immediately following the interruption of long-term ART in infant RMs ^5^. Building on this pre-clinical framework, an investigation into the characterization of HIV-1 reservoirs *in-vivo* with ^64^Cu radiolabeled 3BNC117 mAb (targeting the CD4 binding site of gp120 ^12^) failed to detect HIV-1 Env expression in ART-treated or untreated chronically infected individuals ^6^; however, a year later, another clinical trial with ^89^Zr radiolabeled VRC01 mAb (targeting the CD4 binding site of gp120 ^13^) produced some results of significance in select lymph nodes (LNs), bone marrow, and the gut ^7^. Despite similarities in the biodistributions of the radiotracers and the PLWH groups imaged in both trials, the reasons for the divergent outcomes remain unclear, especially given the higher binding affinity observed for the radiolabeled 3BNC117.

As longer half-life radionuclides can extend the biodistribution time, which in turn allows for lower background and better clearance of nonspecific uptake, in this study, we utilized ^89^Zr radiolabeled intact murine 7D3 and ^89^Zr radiolabeled F(ab’)_2_ fragment of rhesus ITS103.01LS (targeting the CD4 binding site of gp120 ^14^) immunoPET to interrogate the whole-body SIV dissemination *in-vivo* up to 6 days post-radioligand injection and to visualize the localization of SIV virus and viral envelope expressing cells and tissues in chronic and pre-acute infection of adult RMs.

## Results

### In-vitro performance of ^89^Zr-7D3

#### Stability of ^89^Zr-7D3

From radio-HPLC tests, ^89^Zr-7D3 stored in PBS was stable at 4°C for at least 5 days and at 37°C for at least 42h post-radiolabeling (Supplementary Fig. 1A).

#### Binding affinity and binding specificity of ^89^Zr-7D3 to SIV Env expressing cell lines, SIV gp120 coated wells, and primary cells from SIV-infected macaques

^89^Zr-7D3 bound to SIV gp120 protein with a nanomolar affinity (the average equilibrium disassociation constant K_d_ ∼0.35nM from 3 saturation binding assays; Supplementary Fig. 1B). Binding assays with ^89^Zr-7D3 concentration ranging from 0.15nM-6.7nM showed an average binding specificity (expressed as total binding to non-specific binding ratio) to SIV Env expressing cells (activated SIV1C, or MT4 cells infected *in-vitro* with SIVmac251 or SIVmac239-nef-stop) of 7.6 (range 2.6 – 24.7; n=19 assays; 8 assays at 0.15nM, 8 assays at 0.5nM, and 1 assay each at 1nM, 3nM, and 6.7nM concentration). Similarly, the average binding specificity of ^89^Zr-7D3 at concentrations ranging from 0.1nM-7nM to SIV gp120 protein coated on wells was 4.7 (range 3.0 – 6.8; 3 assays each at 0.1, 0.5, 1, 3, 5, and 7nM). However, the binding specificities of ^89^Zr-7D3 to peripheral blood mononuclear cells (PBMC), lymph node mononuclear cells (LNMC), and spleen cells from chronically SIV-infected RMs (Supplementary Table 3) at 0.5nM concentration were similar to the binding specificity of ≈1 observed in negative control cells (cells from uninfected RMs or uninfected MT4 cells).

#### Autoradiography of ^89^Zr-7D3

The autoradiography of LN, spleen, and gut tissue sections from chronically SIV-infected RMs paired with uninfected controls and competed with excess unlabeled 7D3 also did not detect differences in bound ^89^Zr-7D3 (Supplementary Table 4).

### In-vivo performance of ^89^Zr-7D3

#### PET/CT imaging of chronic SIV-infection

After ensuring with radio-HPLC of no formation of immune complexes due to any pre-existing immune response in RMs against the 7D3 mAb or due to other endogenous proteins that may interfere with the antigen binding site of 7D3, we administered ^89^Zr-7D3 intravenously in five chronically SIV-infected RMs and six uninfected controls (Supplementary Table 1). All 11 RMs underwent PET/CT scanning at ∼40h, and 9 of the 11 RMs were scanned again on Day 5-6 post ^89^Zr-7D3 administration. To corroborate *in-vivo* observations, we euthanized one chronically SIV-infected RM and one uninfected control (RM1 and RM8) soon after their ∼40h scan, and another pair (RM4 and RM13) on Day 7 post ^89^Zr-7D3 administration, harvested their organs, removed contents from the bowel, and measured the radioactivity in tissues using a gamma counter. Despite high levels of specificity of binding of ^89^Zr-7D3 to gp120 was observed in *in-vitro* assays, no marked differences of ^89^Zr-7D3 uptake in the PET images were evident visually and semi-quantitatively in any organs, including LNs, spleen, and the gut between the uninfected and the chronic SIV-infected RMs (Fig. 1 and Supplementary Fig. 2). Of note, the radiotracer uptake observed in the colon of the SIV chronically infected RM3, and at a lower intensity in the uninfected control RM10, in the 40h PET scan disappeared at Day 5 (Fig. 1A), suggesting a non-specific intraluminal probe excretion. Similar levels of bowel clearance of the probe were observed in other SIV-infected and uninfected animals. The *ex-vivo* tissue analysis agreed with the *in-vivo* observations (Supplementary Fig. 3A, B).

**Fig. 1.**
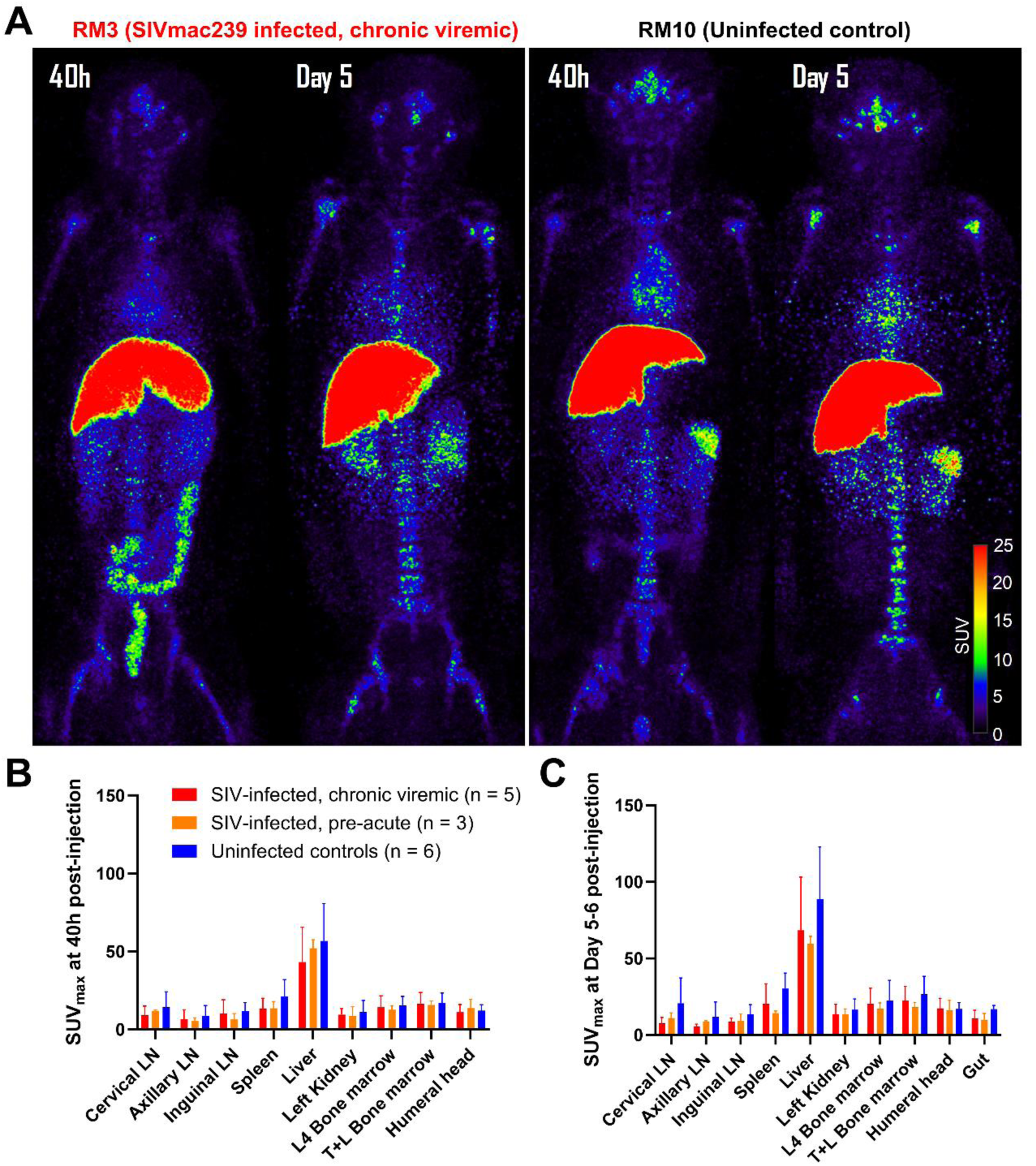
Maximum intensity projection *in-vivo* PET images of a representative chronically SIV- infected viremic rhesus macaque (RM3) and a representative uninfected control (RM10) following administration of ∼1mg mass of ^89^Zr-7D3 (∼0.8mCi of ^89^Zr) and scanned at 40h and Day 5 post- injection **(A)**. Tissue uptakes were converted to RAINBOW color map as shown in the color bar, where red color indicates the high standardized uptake value (SUV). Comparison of maximum SUV (SUV_max_) in tissues among the chronically SIV-infected viremic (red), pre-acutely SIV- infected (orange), and uninfected controls (blue) at 40h (**B**) and Day 5-6 (**C**) post-injection. Both visual and semi-quantitative SUV analysis showed no higher uptake in the SIV-infected RMs compared to the uninfected controls. Because of intraluminal uptake in some animals at 40h, the gut uptake is not plotted. Plots are mean values and error bars are standard deviation.

At any time point measured post-injection, we observed no significant differences in the standardized uptake value (SUV) of ^89^Zr-7D3 in the peripheral blood and plasma between SIV-infected and uninfected (P = NS; Supplementary Fig. 4B, C). Using the radioactive plasma obtained at ∼13h post-injection (mean±standard deviation estimated ^89^Zr-7D3 concentration in the plasma was 7.1 ±2.9nM (n=11)), we performed a binding assay in a subgroup of animals (4 chronically infected and 3 uninfected). ^89^Zr-7D3 in the plasma retained its binding specificity to SIV gp120 only in the uninfected controls but not in the chronically infected RMs in which >97% reduction in binding to gp120 was observed. Even in the binding assay with radioactive plasma obtained at earlier times (1h, 2h, 4h, and 6h post-injection) when ^89^Zr-7D3 concentration was much higher (calculated to be ∼18nM at 1h post-injection, n=3 (2 chronically infected and 1 uninfected)), we observed a near full loss of ^89^Zr-7D3 binding to gp120 in the plasma of the chronically infected RMs at all time points, but not in the uninfected plasma (data not shown).

#### In-vivo Stability of ^89^Zr-7D3

Radio-HPLC of the plasma obtained at ∼13h (data not shown) and ∼40h post ^89^Zr-7D3 injection in all 11 RMs (5 chronically SIV-infected and 6 uninfected RMs) confirmed near 100% stability of ^89^Zr-7D3 *in-vivo,* and we detected no formation of high molecular weight complexes nor breakdown to small molecules (Supplementary Fig. 1D), suggesting that the binding loss of ^89^Zr- 7D3 probe in the plasma of the chronically SIV-infected RMs was likely due to high concentrations of endogenous proteins competing for the 7D3 binding site.

### Development of endogenous proteins in SIV-infected macaques that compete for the 7D3 binding site

To explore the rapid loss of binding of ^89^Zr-7D3 probe in the plasma of the chronically SIV-infected RMs, we tested the *in-vitro* binding of ^89^Zr-7D3 to SIV Env expressing cells in the presence of uninfected or chronic plasma (Supplementary Table 5). The 7D3 binding in the presence of chronic plasma was fully abrogated, with the binding similar to non-specific binding wells where Env+ cells were pre-incubated with ∼3-4 Log_10_ higher concentration of the unlabeled 7D3. We then tested the emergence of binding inhibition in five RMs intravenously inoculated with 200 TCID_50_ of SIVmac239-nef-stop and followed longitudinally ^15^. Binding assay in the presence of plasma revealed no loss of 7D3 binding to SIV Env expressing cells at Day 7 post-inoculation (p.i.), 0- 51% loss at Day 14 p.i., and near full binding abrogation by week 5 p.i (Fig. 2A). Radio-HPLC of the incubated ^89^Zr-7D3 + plasma mixture revealed no formation of high molecular weight immune complexes and >90% of the radioactivity of the injected ^89^Zr-7D3 + plasma mixture eluted with a retention time identical to that of ^89^Zr-7D3, suggesting that the binding inhibition was due to endogenous proteins, likely antibodies, in the chronic plasma competing for the 7D3 binding site and not due to the blockade of the 7D3 antigen binding site (Supplementary Fig. 1E). A similar gradual 7D3 binding loss was also observed during SIVmac251- and SIVmac239-infection where endogenous proteins competing for 7D3 binding site were either absent or below the assay detection levels in the plasma for at least until 10 days post-inoculation (Fig. 2B, C). Though no loss of 7D3 binding to SIV Env expressing cells was observed in the presence of pre-acute plasma, the binding specificity of ^89^Zr-7D3 to Day 7 and Day 10 PBMC was ≈1 (Supplementary Table 3).

**Fig. 2.**
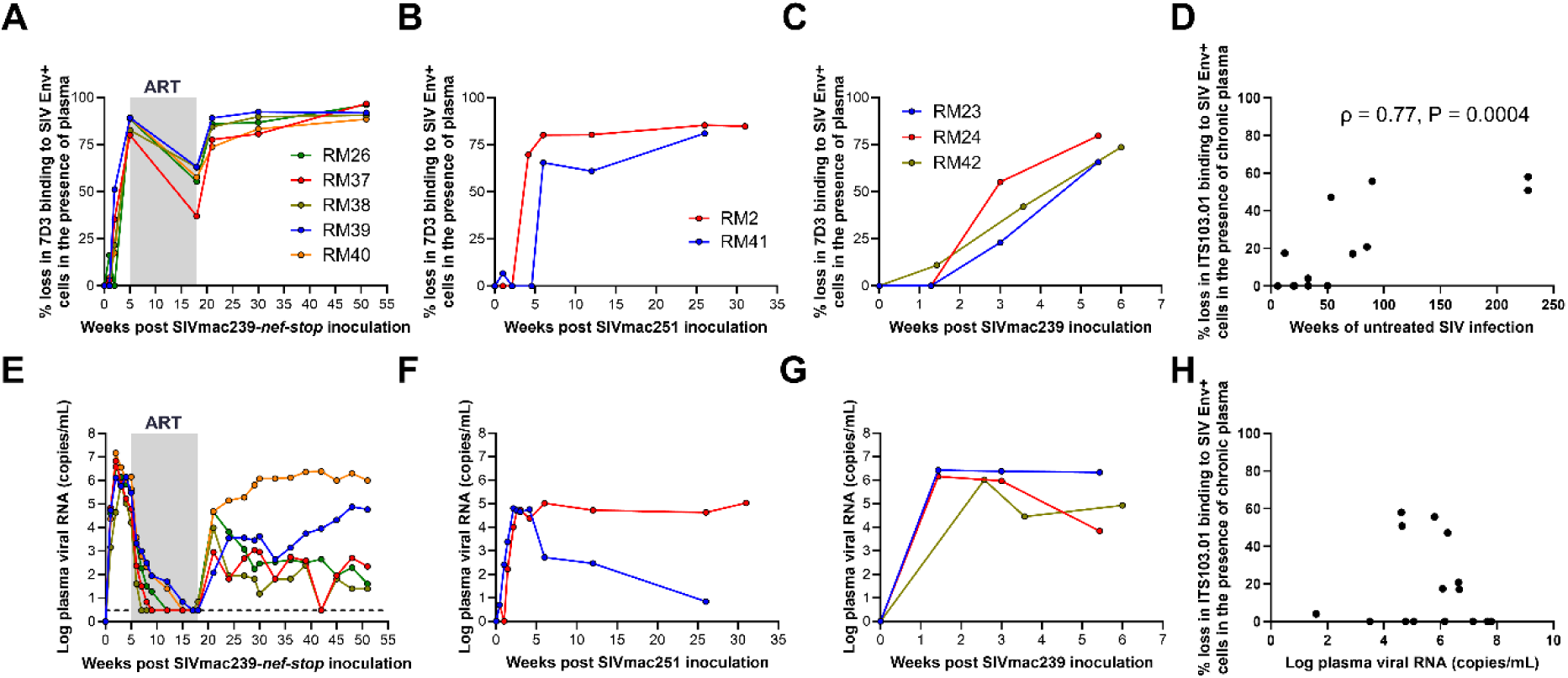
Percent loss of 7D3 binding to SIV Env+ cells in the presence of plasma from 5 rhesus macaques intravenously inoculated with 200 TCID_50_ SIVmac239-nef-stop **(A)**, 2 rhesus macaques intravenously inoculated with 300 TCID_50_ SIVmac251 **(B)**, 3 rhesus macaques intravenously inoculated with 1000 TCID_50_ SIVmac239 **(C)**, and their respective plasma viral RNA (**E, F, G**). Percent loss of ITS103.01LS-F(ab’)_2_ binding to SIV Env+ cells in the presence of chronic plasma from 15 rhesus macaques infected with either SIVmac239 or SIVmac239-nef-stop and either antiretroviral therapy naïve or off antiretroviral therapy (**D, H**). The horizontal dashed line in panel E indicates the minimum detection threshold of 3 copies/mL. In panel D, ρ is the Spearman rank correlation coefficient and P is one-sided significance.

After five weeks of SIVmac239-nef-stop infection, the five RMs initiated a 13-week course of daily antiretroviral (ART) ^15^. At the end of 13 weeks of effective ART, we observed a reduced loss in 7D3 binding to Env+ cells in the presence of ART-suppressed plasma (plasma viral load (PVL) <10 copies/mL) (Fig. 2A, E). Three weeks following ART interruption, a near-complete abrogation of 7D3 binding, and during the chronic phase of infection, a complete 7D3 binding abrogation was observed in all five SIVmac239-nef-stop-infection RMs (Fig. 2A). However, in a group of four SIVmac251 infected macaques that were rectally challenged with 1000 TCID_50_ and chronically infected for over 1 year before initiating ART, complete inhibition of 7D3 binding to Env+ cells was observed in the presence of ART-suppressed plasma (on ART for >6 months (range 32 to 78 weeks), PVL <50 copies/mL at least for the last 6 months) (data not shown).

### PET/CT imaging of pre-acute SIV-infection with ^89^Zr-7D3

As the failure of ^89^Zr-7D3 to highlight *in-vivo* differences of SIV Env expression between chronically infected and uninfected controls could in part be due to excess levels of endogenous proteins competing for the gp120 binding site of 7D3, we hypothesized that the potential to observe *in-vivo* SIV Env expression could be during pre-acute infection before an antibody response to gp120 was mounted. So, we intravenously inoculated three RMs with SIVmac239 (Supplementary Table 1). After screening for any pre-existing immune response against the 7D3 mAb and also confirming the absence of interfering and competing endogenous proteins in the plasma on Day 7 of SIV-infection, we administered ^89^Zr-7D3 on Day 8 of SIV-infection and performed PET/CT imaging at ∼40h (Day 10), optionally at ∼90h (Day 12), and ∼108-132h (Day 13-14) post ^89^Zr- 7D3 administration. Radio-HPLC of the plasma obtained at ∼10h (data not shown) and ∼40h post ^89^Zr-7D3 injection confirmed near 100% stability of ^89^Zr-7D3 *in-vivo* (Supplementary Fig. 1D). Both qualitative and semi-quantitative SUV analyses of various lymphoid tissues and the gut failed to show higher uptake in the SIV-infected compared to the uninfected controls. (Supplementary Fig. 2, 5, and Fig. 1B, 1C).

Since the levels of viral RNA in lymphoid tissues were shown to be similar between Day 7-10 post-infection and the chronic phase of SIV-infection ^16–18^, we combined the five chronic and three pre-acutely infected RMs and compared the uptake with the six uninfected controls. We observed no statistically significant differences in the radiotracer uptake between SIV-infected and uninfected controls (P = NS).

### Immunogenicity of ^89^Zr-7D3

We tested the immunogenicity of the murine 7D3 mAb probe 5-11 weeks after exposure in 9 of 14 RMs (2 chronic, 3 pre-acute, and 4 uninfected). Though HPLC showed no formation of high molecular weight immune complexes in all 4 uninfected RMs and the 2 chronic RMs after the exposure, 2 of 3 acutely SIV-infected macaques developed an immune response against the 7D3 mAb.

Though comparably viremic, the pre-acute infection may not translate to similar gp120 concentrations present in the tissues of chronically infected animals. To image chronic infection, we found ITS103.01LS to be a promising mAb with good binding affinity and binding specificity to SIV gp120, and also less unaffected by the barrier of competing endogenous proteins consistently detected for 7D3.

### In-vitro performance of ^89^Zr-ITS103.01LS-F(ab’)_2_

#### Stability of ^89^Zr-ITS103.01LS-F(ab’)_2_

^89^Zr-ITS103.01LS-F(ab’)_2_ stored in PBS was stable at 4°C for at least 6 days and at 37°C for at least 48h post-radiolabeling as assessed by radio-HPLC (Supplementary Fig. 1F).

#### Binding affinity and binding specificity of ^89^Zr-ITS103.01LS-F(ab’)_2_ to SIV Env expressing cell lines, SIV gp120 coated wells, and primary cells from SIV-infected macaques

Saturation binding assays of ^89^Zr-ITS103.01LS-F(ab’)_2_ showed a nanomolar affinity to gp120 (K_d_ ∼0.87nM, n=4 assays; Supplementary Fig. 1C). In binding assays at 0.5nM concentration of ^89^Zr- ITS103.01LS-F(ab’)_2_, we observed an average binding specificity of 6.1 (range 3.2 – 8.7; n=9 assays) to SIV Env expressing cells (MT4 cells infected *in-vitro* with SIVmac239-nef-stop) and an average binding specificity of 5.4 (range 4.8 – 6.0; n=5 assays) to SIV gp120 coated on wells. Yet, as observed with 7D3, the binding specificities of ^89^Zr-ITS103.01LS-F(ab’)_2_ to PBMC, LNMC, and spleen cells of chronically SIV-infected RMs (Table S6) at 0.5nM concentration were similar to the binding specificity to cells from uninfected controls or uninfected MT4 cells of ≈1.

#### Autoradiography of ^89^Zr-ITS103.01LS-F(ab’)_2_

No differences in the binding specificity of ^89^Zr-ITS103.01LS-F(ab’)_2_ were noted on the autoradiography of LN, spleen, and gut tissue sections from chronically SIV-infected RMs compared with uninfected controls or when competed with excess unlabeled ITS103.01LS (Supplementary Table 7).

### Endogenous proteins in SIV-infected macaques competing for the ITS103.01LS-F(ab’)_2_ binding site

Though we still observed some loss of ITS103.01LS-F(ab’)_2_ binding to SIV Env expressing cells in the presence of some chronic plasma samples due to competing endogenous proteins, the loss was more variable and significantly lower than 7D3 binding loss (% loss = 0-58%, average loss = 18%, n=15, Log_10_ PVL range = 1.6-7.8 copies/mL) with some animals showing no loss, and as expected, the loss strongly correlated with the length of untreated SIV-infection (ρ = 0.77, P < 0.001, Fig. 2D).

### In-vivo performance of ^89^Zr-ITS103.01LS-F(ab’)_2_

#### PET/CT imaging of chronic SIV-infection

We administered ^89^Zr-ITS103.01LS-F(ab’)_2_ in five SIVmac239 chronically infected RMs and five uninfected controls (Supplementary Table 2) and performed PET/CT scans at ∼48h and Day 5-6 post-injection. In two RMs (RM10 and RM16) we observed the formation of high molecular weight complexes without loss in binding to SIV Env expressing cells, and in two other RMs (RM3 and RM15) we detected endogenous proteins competing for the binding site of ITS103.01 mAb resulting in ∼20% loss of binding of ^89^Zr-ITS103.01LS-F(ab’)_2_ to SIV Env expressing cells. The plasmas of all other animals showed no changes in the molecular weight of the probe and no loss of binding of the probe to SIV Env expressing cells. Both qualitative and semi-quantitative PET SUV analysis of various lymphoid tissues and the gut failed to show higher uptake in the chronic SIV-infected RMs compared to the uninfected controls (Fig. 3 and Supplementary Fig. 6). The probe uptake observed in the gut of some RMs at 48h cleared by Day 5 and is consistent with intraluminal uptake (data not shown). Two RMs (RM3 and RM17) were euthanized soon after their Day 5 imaging and organs were harvested to corroborate *in-vivo* observations. The *ex-vivo* tissue analysis in a gamma counter agreed with the *in-vivo* images (Supplementary Fig. 3C).

**Fig. 3.**
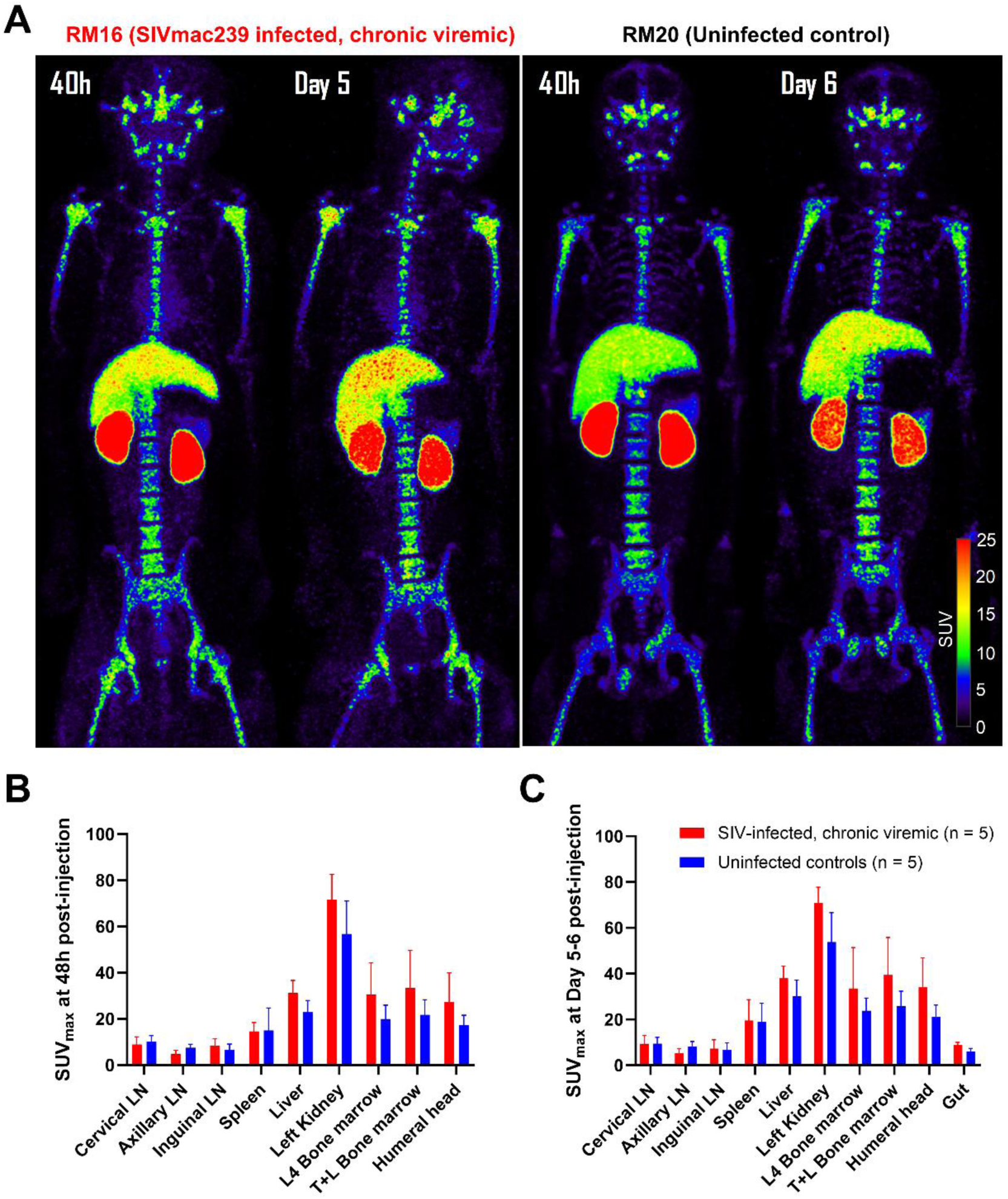
Maximum intensity projection *in-vivo* PET images of a representative chronically SIV- infected viremic rhesus macaque (RM16) and a representative uninfected control (RM20) following administration of ∼1mg mass of ^89^Zr-ITS103.01LS-F(ab’)_2_ (∼2mCi of ^89^Zr) and scanned at 48h and Day 5-6 post-injection **(A)**. Tissue uptakes were converted to RAINBOW color map as shown in the color bar, where red color indicates the high standardized uptake value (SUV). Comparison of maximum SUV (SUV_max_) in tissues among the chronically SIV-infected viremic (red) and uninfected controls (blue) at 48h (**B**) and Day 5-6 (**C**) post-injection. Both visual and semi-quantitative SUV analysis showed no higher uptake in the SIV-infected RMs compared to the uninfected controls. Because of intraluminal uptake in some animals at 48h, the gut uptake is not plotted. Plots are mean values and error bars are standard deviation.

#### In-vivo stability of ^89^Zr-ITS103.01LS-F(ab’)_2_

We obtained plasma at ∼9h post ^89^Zr-ITS103.01LS-F(ab’)_2_ injection in 8 of the 10 RMs (4 chronically infected and 4 uninfected). Radio-HPLC of that plasma confirmed near 100% stability of the radiotracer *in-vivo* (Supplementary Fig. 1G), and a binding assay confirmed the probe in the plasma of all 8 RMs (mean (±standard deviation) estimated ^89^Zr-ITS103.01LS-F(ab’)_2_ concentration was 10.0 (±1.5)nM (n=8)) retained its binding specificity to SIV Env expressing cells. As the ^89^Zr-ITS103.01LS-F(ab’)_2_ tracer cleared faster than ^89^Zr-7D3 (Supplementary Fig. 4E, F), we could not run *in-vivo* stability analysis with HPLC at later time points. However, ^89^Zr- ITS103.01LS-F(ab’)_2_ incubated with RM plasma *in-vitro* at 37°C was near 100% stable for at least 48h.

### Immunogenicity of ^89^Zr-ITS103.01LS-F(ab’)_2_

We tested the immunogenicity of the rhesus ^89^Zr-ITS103.01LS-F(ab’)_2_ mAb probe 5-10 weeks after exposure in 5 of 10 RMs (3 chronic and 2 uninfected). Radio-HPLC assay showed no development of immune response against the ITS103.01LS-F(ab’)_2_ mAb in all 5 RMs tested, nor were there changes in immunoreactivity.

## Discussion

Two pre-clinical studies by Santangelo et al.^1,2^ using 7D3 mAb, targeting gp120, claimed feasibility to non-invasively detect, localize, and quantify viral sites in chronic and acute SIV infection, as well as the residual virus in antiretroviral-treated aviremic RMs (when the viral RNA in lymphoid tissues is reduced by ∼4 Log_10_). Therefore, we selected the same 7D3 mAb, radiolabeled it with ^89^Zr, and observed an *in-vitro* binding specificity of ∼8 to SIV cell lines. However, the binding specificity to primary cells (PBMC, LNMC, and spleen cells; freshly isolated and/or cryopreserved) from SIV-infected RMs was ∼1, indicating a lack of specific binding. In contrast, with ^64^Cu-DOTA-7D3-PEG, Santangelo et al.^1^ reported an *in-vitro* binding specificity of ∼3 using cell lines, and ∼2-3 to the spleen and LN cells from SIV-infected RMs. Of note, Santangelo et al. used cryopreserved (but not freshly isolated) cells from fewer animals and without cold-competition assay, all potential factors contributing to the discrepancy with their study.

Since not all gp120 in tissue is expressed on the primary cells, we also performed autoradiography, an *ex-vivo* validating technique that often yields data with robust predictive power regarding the feasibility of imaging a target *in-vivo*, which was lacking in all previous studies claiming the feasibility of imaging SIV\SHIV\HIV viral replication *in-vivo*. Autoradiographic analyses of ^89^Zr- 7D3 consistently showed no differences between total binding and non-specific binding of tissue sections from several lymphoid organs of SIV-infected animals or compared to uninfected animals.

Consistent with the *ex-vivo* measurements, we found no discernible differences in ^89^Zr-7D3 uptake in any organs between SIV-infected and uninfected animals *in-vivo*. Radio-HPLC analyses of the radiotracer circulating in the plasmas of imaged animals confirmed high stability of ^89^Zr-7D3 *in-vivo*. We detected evidence of endogenous interference (likely antibodies) competing for the binding site of 7D3 in SIV chronically infected RMs at concentrations sufficient to fully abrogate binding of the ^89^Zr-7D3 to Env+ cell lines, i.e., cells with gp120 expressed at levels much higher than primary cells. Santangelo et al.^1^ also observed interference of 7D3 binding to Env+ cells in the presence of post-infection serum; however, comparing the relative titers of endogenous antibodies between the two studies is not feasible due to the utilization of different assays (flow cytometry ^1^ vs. radiotracer binding assay in our study). Thus, to explain the discrepant outcomes, one may be tempted to speculate that our chronic SIV RMs had developed endogenous antibodies at levels higher than those elicited in the animals imaged in all the earlier 7D3 imaging studies ^1–4^.

The pre-acutely infected animals imaged in our study with ^89^Zr-7D3 (with PVL >10^6^ SIV-RNA copies/ml), on the other hand, dismiss competing antibodies as a potential explanation for our inability to image the virus *in-vivo*, because these hosts had not yet had sufficient time, at the time of imaging, to develop such endogenous interference as confirmed through *in-vitro* plasma binding assays. Although we had not collected tissues at the time of imaging to confirm that sufficiently high levels of viral dissemination in lymphoid organs had been reached, LNs biopsied at week one post-SIVmac infection of both cynomolgus and rhesus macaques have been reported to achieve CAVL levels similar to those found in SIV chronically infected animals ^16–18^. Nevertheless, it is important to recognize that the gp120 molarity in tissues of SIV- or HIV-infected hosts as well as its association with CAVL-RNA\DNA levels in the same tissues have been minimally investigated to date ^19^. To account for all these potentially confounding variables, we next evaluated the performance of a second anti-gp120 radiotracer, ^89^Zr-ITS103.01LS-F(ab’)_2_, after discovering evidence of significantly lower (or absent) endogenous antibodies in the plasmas of SIV chronically infected animals competing for its binding to gp120, possibly due to differential immunogenicity of the CCR5 and CD4 binding sites of the gp120. ^89^Zr-ITS103.01LS-F(ab’)_2_ also showed high specificity to Env+ cells (∼6) as well as binding affinity (∼1nM) to gp120 similar to those observed for ^89^Zr-7D3. Yet, like ^89^Zr-7D3, ^89^Zr-ITS103.01LS-F(ab’)_2_ also failed to detect evidence of specific binding to primary cells, autoradiography tissue sections, as well as *in-vivo* in SIV-chronically infected RMs, including in animals with absent levels of endogenous antibodies competing for the same binding site. Both ^89^Zr-7D3 and ^89^Zr-ITS103.01LS-F(ab’)_2_ datasets collectively rule out the possibility that the lack of evidence for specific binding in our study is attributable to higher levels of endogenous antibodies in our infected animals interfering with the binding of the probe to gp120, suggesting that the binding potential (BP; the product of the ligand’s affinity and the target’s molarity ^20^) of these probes is too low for imaging gp120 levels in tissues *in-vivo*. Indeed, the only evidence of specific binding in our study was observed in gp120- expressing cell lines. Of note, the cell-associated viral load (CAVL) measured in SIVmac-infected MT4 cell lines with binding specificity of 5.1-7.9 was ∼10^10^-10^11^ copies/million cells (data not shown), hence ∼5 Log_10_ higher than the typical SIV-RNA levels observed in the LNs or rectal tissues of highly viremic chronically SIV-infected RMs ^15^, and ∼8-9 Log_10_ higher than the residual viral replication in ART-treated, PVL-suppressed, SIV-infected RMs ^15^.

Despite utilizing similar mAb mass, injected radioactivity amount, and animal weights, the SUV levels in the two intact 7D3 studies we sought to replicate appear to be dramatically low (up to ∼1 Log_10_ lower compared to our study) ^1,2^. While PEGylation could, in principle, affect biodistribution by increasing molecular size, as well as by decreasing non-specific uptake ^21^, this modification, in our opinion, would not be able to explain the lack of differences in SUV levels between uninfected and infected tissues observed in all our assays (*in-vitro*, *ex-vivo*, and *in-vivo*), nor the lower SUV levels found in their studies. Although the stability, radiochemical purity, and whole-body retention of the radiotracer were not reported in the previous pre-clinical studies, in our study, ^89^Zr- 7D3 was stable *in-vivo* and *in-vitro*, with high whole-body retention (average ∼72%) at ∼40h post-injection (Supplementary Fig. 4A), consistent with the biodistributions of other radiolabeled intact antibodies ^22,23^. Furthermore, the radiometal (^89^Zr vs. ^64^Cu) does not appear to account for the observed differences in SUV levels in large organs (e.g. kidneys and liver) between our study and the previous intact 7D3 studies ^1,2^ as the biodistribution of the intact 7D3 mAb radiolabeled with ^64^Cu (^64^Cu-DOTA-7D3) and PET imaged at 36h post-injection in our program showed whole-body retention and SUVs in organs similar to those observed with ^89^Zr-7D3 (data not shown).

Perplexingly, Santangelo et al.^1^ demonstrated evidence of higher SUV levels in the SIV-infected RMs compared to the uninfected RMs not only in LNs and the spleen, the tissues known to harbor high levels of viral dissemination, but also in other organs such as the kidneys, liver, and heart, which typically contain much lower RNA levels ^24^. The >2-fold higher uptake of the probe in the heart (blood pool) of infected RMs, indicative of the input function, suggests indeed a non-specific nature of the increase in SUV levels found in other organs, e.g. axillary LNs for which less than 2-fold increase in SUV levels were reported, suggesting a decrease in relative uptake compared to the blood (SUV_LN_/SUV_heart_).

Adopting the same immunoPET methodology, in late 2022, Samer et al.^3^ claimed the feasibility of imaging LRA-induced viral reactivation in tissues of long-term antiretroviral-treated aviremic SIV-infected RMs using the non-pegylated F(ab’)_2_ fragment of the primatized 7D3 radiolabeled with ^64^Cu, although no *in-vitro* binding data in primary cells or tissue sections were provided to corroborate their observations, nor included an uninfected control arm. Increased SUV levels were reported in axillary LNs and the gut following one cycle of the LRA administration for one or two weeks but without concomitant increases in heart SUV levels. However, we believe that those asymmetrical SUV increases observed in axillary LNs ipsilateral to the injection site combined with the lack of increase in uptake in other clusters of LNs (e.g. the inguinal LNs which showed increases in viral RNA levels following the LRA administration comparable to axillary LNs) likely indicate fully non-specific uptake due to accidental infiltration of the probe into subcutaneous space observed in the imaging time points post-LRA administration but not in the baseline imaging time points of several RMs (four out of the five animals for which increase in nodal SUV levels were reported). Subcutaneous administration of radiolabeled antibodies is known to result in non-specific LN uptake ^25^. The gut uptake, moreover, appears consistent with intraluminal excretion, also observed with other radiolabeled intact mAbs or mAb fragments ^7,26^, which seems included in their studies given how their regions of interest were drawn. Of note, similar high levels of probe uptake in the gut were observed in some animals in our study at 40-48h post-injection, which dramatically cleared by Day 5 post-injection (a phenomenon that we were able to observe in our study due to the longer half-life of the ^89^Zr compared to ^64^Cu), again indicating a non-specific nature of such uptake.

More recently (Kim et al.^4^), the same team conducted the same immunoPET imaging during multiple cycles of 2-week on/2-week off LRA administration at higher doses while the RMs were on suppressive ART. A notable rise in LN and gut SUV levels during the third cycle coincided with increased blood pool (heart) and liver SUV uptake. In conjunction with the earlier NHP studies by Santangelo et al. where increased LN uptake coincided with increased heart activity, it suggests the latter is necessary for observing increased LN SUV levels with their imaging system. This holds especially true if we consider the LN increase reported in Samer et al. occurring without a corresponding heart increase was due to tracer extravasation. While Kim et al.^4^ do not dismiss nonspecific uptake as an explanation for the observed increase in blood pool activity (e.g. due to galunisertib-induced alteration in probe pharmacokinetics or probe-antigen plasma kinetics), they also suggest that the observed increase in heart SUV during the third cycle could be due to changes in viral antigen concentrations, consistent with the feasibility of imaging SIV *in-vivo* claimed in the previous three NHP immunoPET studies ^1–3^. Unfortunately, Kim et al. ^4^ did not include a control group of uninfected RMs to exclude that the induced changes in SUV levels observed in their SIV-infected animals could be explained by non-specific alteration in antibody biodistribution caused by the drug regimen, hence unrelated to SIV presence. Of note, if the addition of a control group shows that, post-galunisertib administration, changes in SUV uptakes are observed only in SIV-infected animals, it may still be the result of non-specific alteration in antibody-gp120 complex kinetics, as implied by the same authors ^4^, hence again unrelated to gp120 molarity in tissues. Though, to our knowledge, SIV gp120 levels in whole blood have not been reported in the literature, HIV-1 and SHIV studies suggest gp120 levels of ∼1 ng/mL of plasma during acute and early stages of infection ^27,28^. Assuming all gp120 in the blood is available for binding (and not occupied by endogenous anti-gp120 antibodies) and using the reported probe’s specific activity in Kim et al. ^4^, we estimate that <0.1% of blood pool activity may be specific binding. Moreover, we do not see evidence of the formation of probe-antigen complexes in the plasmas of any of our SIV-infected animals (both *in-vitro* and *in-vivo*) based on radio-HPLC analyses; nor do we see interference in binding to gp120 expressing cell lines from ^89^Zr- ITS103.01LS-F(ab’)_2_ incubated in plasmas of highly viremic RMs. Whether the LRA induces shedding of gp120 in the plasma to achieve levels much higher than what is expected in an infected animal matched for plasma viremia and sufficient to significantly occupy the probe is nevertheless unknown; yet, the latter would again indicate a change in biodistribution of the probe (bound to the gp120) fully non-specific, hence unrelated to changes in gp120 molarity in tissues. And again, as reported in their prior study ^3^, the increase in probe uptake was observed only in the axillary LN cluster but not in the spleen. It is also crucial to note that the probe uptake in larger organs (such as the liver, kidney, and spleen) after the third cycle remained notably lower compared to both radiotracers in our study. This necessitated the use of different SUV display scales ((0-1.5) in their studies ^3,4^ and (0-25) in our study, suggesting that the majority of the ^64^Cu-p7D3-F(ab’)_2_ radiotracer was rapidly cleared from the body within 24 hours of intravenous infusion, in contrast with the ^89^Zr-ITS103.01LS-F(ab’)_2_ radiotracer in this study where on average ∼74% remained in the body at ∼48h post-injection (Supplementary Fig. 4D). Notably, hepatic and renal SUV levels in our study are similar to those reported for other radiolabeled F(ab’)_2_ fragments (with ^99m^Tc, ^89^Zr, or ^64^Cu) at similar time points post-radioligand injection in rhesus macaques ^29,30^ and humans (with ^111^In) ^31^, as well as for ITS103.01LS-F(ab’)_2_ radiolabeled with ^64^Cu (^64^Cu-DOTA-ITS103.01LS- F(ab’)_2_) and PET imaged at 24h post-injection in our program (data not shown).

In summary, given that the increase in SUV levels in tissues has been observed in these four NHP immunoPET studies ^1–4^ only when there is a concomitant increase in SUV levels of the heart, that none of these studies have documented evidence of specific uptake *ex-vivo* in tissues, and that the evidence of specific uptake in primary cells was reported only in the first publication with a limited number of samples and without competition assay ^1^, we believe that the feasibility of imaging SIV *in-vivo* with immunoPET using anti-Env antibodies\fragments with nanomolar affinities has not been demonstrated to date and the discrepancy in the biodistribution between our study and earlier NHP imaging studies remains unexplained. Therefore, it is crucial, given the state-of-the-art research in this novel area of imaging science, to contextualize the efforts of the past decade in demonstrating the feasibility of imaging SIV/HIV viral replication *in-viv*o using non-invasive nuclear medicine technologies and compare with milestones achieved in other areas of imaging viruses *in-vivo*.

Two decades ago, when imaging science initiatives began harnessing nuclear medicine technologies for *in-vivo* viral replication imaging ^32^, the consensus in the field was that only the herpes simplex virus (HSV) could be imaged *in-vivo* with a radiolabeled tracer. This was achieved by exploiting a mechanism of signal amplification through viral thymidine kinase (TK), which phosphorylates and traps the probe within infected cells. Imaging viruses that lack such enzymes for signal amplification was anticipated to rely primarily on quantitative aspects of ligand-receptor mechanisms, such as radiolabeling antibodies to specific targets on virions or infected cells. In general, the feasibility of imaging a target with this mechanism primarily depends on BP, which must be sufficiently high for successful *in-vivo* imaging of the virus. Additionally, for a given BP, the higher the association rate, the greater the feasibility of imaging the target at earlier timepoints post-injection. However, other factors beyond BP, such as biodistribution, may prevent the imaging system from visualizing the viral target *in-vivo*, emphasizing the importance of *ex-vivo* autoradiography studies to predict success in imaging viruses *in-vivo* ^32^. To clarify, the presence of specific uptake in tissue sections from infected hosts suggests the potential for detecting the virus *in vivo*. Conversely, the absence of such evidence in autoradiography tissue sections indicates a considerable setback in the feasibility of *in-vivo* viral imaging.

The lack of specific uptake observed in our study in primary cells and tissues obtained from SIV-infected hosts with two different probes exhibiting nanomolar binding affinities to two different sites of the gp120 suggests that the BP of such probes may not be sufficiently high to succeed in imaging SIV *in-vivo*. If our negative findings are validated, alternative imaging approaches should be interrogated involving higher affinity ligands for gp120, although to our knowledge no such ligands are yet reported in the literature, or other viral targets such as internal viral proteins like capsid, which are the most abundant, or proteases, especially considering their reported picomolar affinity ^32^. Additionally, examining mechanisms of signal amplification, as proposed by Bhaumik et al.^33^ warrants investigation in preclinical settings.

Yet, inspired by the successes claimed in the earlier SIV NHP publications, two clinical imaging studies utilizing non-pegylated anti-Env bNAbs, ^89^Zr-VRC01 (Beckford-Vera et al. ^7^) and ^64^Cu- 3BNC117 (McMahon et al.^6^), undertook to non-invasively image HIV-1 but reached conflicting conclusions. As expected, the biodistributions of the two radiotracers appeared remarkably similar. While the shorter half-life of ^64^Cu allowed McMahon et al. to image patients for up to 48h post-radiotracer injection, Beckford-Vera et al. imaged patients up to 72h post-injection. Notably, the ^89^Zr-VRC01 study saw a clear trend for higher uptake in the LNs and the gut of HIV-1 infected subjects already at Day 0 and 1, hence ruling out the longer half-life of the ^89^Zr as an explanation for the contrasting conclusions. Additionally, the affinity of ^64^Cu-3BNC117 appeared to be ∼2- fold higher than that of ^89^Zr-VRC01, suggesting that the BP should be higher for the former, given that the two studies enrolled similar groups of HIV-1 infected viremic individuals. Moreover, it is important to acknowledge that the absolute increase in SUV levels observed in the LNs and gut in the ^89^Zr-VRC01 study is only ∼10-20% of the SUV levels in the heart (blood pool), hence close to background levels as can be clearly appreciated by their maximum intensity projection images, and no significant differences in spleen uptake was observed.

A crucial difference in the quantification of the two clinical studies is that ^64^Cu-3BNC117 used the standardized uptake value (SUV) as an operator, similar to the monkey imaging studies, while ^89^Zr-VRC01 used SUV normalized on the aortic outflow tract (the radioactivity in the blood), i.e., tissue to blood ratio. This different measure could lead to varying interpretations of the data, especially if the blood pool activity is lower in HIV-1 infected individuals, as the published images in both papers seem to highlight, resulting in higher ratios. This issue is particularly important considering the pharmacokinetics of these two bNAbs when administered at pharmacologic doses have shown some evidence of faster clearance in HIV-1 infected individuals ^34,35^, a phenomenon with an unknown mechanism but that appears to pose theoretical challenges when explained solely by sequestration due to binding to gp120 receptors. The authors of the ^89^Zr-VRC01 study do not rule out indeed the possibility that increases in relative SUV observed in LNs and the gut of HIV- 1 infected individuals (both untreated and ART-treated with suppressed viral load) may indicate non-specific uptake in areas of heightened tissue inflammation with concurrent elevation in Fc receptor expression. In theory, this phenomenon, alongside other indirect effects of viral-induced pathogenesis ^36^, could explain the positive statistically significant associations between probe uptakes and viral dissemination levels observed in some lymphoid organs of Beckford-Vera et al. (with n=5 individuals) ^7^ and Samer et al. (with n=7 macaques) ^3^, although the latter contrasts with the lack of differences in SUV levels between infected and uninfected hosts observed in our study. Furthermore, the observation that such signal increase (attributed to an increase in viral replication) is evident in LNs but not in the spleen in the Beckford et al. study as well as in Samer et al. ^3^ and Kim et al. ^4^ studies suggest, in our view, a potential lack of robustness in the imaging systems under scrutiny because the levels of viral replication and/or gp120 (per unit volume of tissue) are known to be similar between LNs and the spleen ^19,24^. Given the partial volume effect of PET cameras, one would anticipate a higher signal in the spleen rather than in the LNs, given their size differences.

Inspired by the first SIV NHP imaging study ^1^, immunoPET methodologies are being adapted to image SARS-CoV-2 infected NHPs ^37^. We anticipate that the outcomes of these investigations will be influenced by quantitative considerations regarding the relative BP of the proposed radiolabeled probes. This involves assessing probe binding affinities and target densities of SIV-gp120, HIV- gp120, and SARS-CoV-2 spike protein in the imaged tissues. For example, the affinities of the anti-HIV-gp120 bNAbs ^89^Zr-VRC01 and ^64^Cu-3BNC117 mentioned earlier are ∼6-15 fold lower and ∼2-6 fold lower compared to the affinities of ^89^Zr-7D3 and ^89^Zr-ITS103.01LS-F(ab’)_2_, respectively. This suggests that assuming similar concentrations of gp120 in SIV and HIV-1 chronically infected hosts with high plasma viremia, the NHP model would likely provide a more sensitive imaging system than imaging the HIV. Conversely, a lack of reproducibility in imaging SIV *in-vivo* has direct theoretical implications for the feasibility of imaging HIV *in-vivo*.

In conclusion, further investigation and consideration of various factors are essential to fully comprehend the implications of the findings from both the preclinical and clinical imaging studies of lentiviral replication discussed above and challenged by our observations.

## Methods

### Animals

Forty-seven juvenile to adult Indian rhesus macaques (RMs) (*Macaca mulatta*) were used in the study following the National Institute of Allergy and Infectious Diseases (NIAID) Animal Care and Use Committee approved protocol. Of the 47 RMs, 20 RMs (designated RM1-RM20) were used for *in-vivo* PET/CT imaging with ^89^Zr-7D3 or ^89^Zr-ITS103.01LS-F(ab’)_2_ as outlined in Tables S1-S2 and some were also used as a source of plasma, cells, or tissue. The remaining 27 RMs (designated RM21-RM47) served exclusively as sources of plasma, primary cells, or tissue for *in-vitro* studies (Tables S3-S7).

The daily antiretroviral therapy (ART) regimen consisted of three antiretroviral drugs comprising two reverse transcriptase inhibitors and one integrase inhibitor: Tenofovir (PMPA, 20mg/kg, Gilead) and Emtricitabine (FTC, 30mg/kg, Gilead), each injected subcutaneously once a day in the scapular area of the back, and either Raltegravir (20mg/kg, Merck) given twice a day mixed in food, or L-870812 (20mg/kg (max 100mg), Merck) given once a day mixed in food.

All procedures were performed at the National Institutes of Health (NIH) Animal Center. Animals’ health was monitored daily and are cared for according to the *Guide for the Care and Use of Laboratory Animals*, 8^th^ Ed., Animal Welfare Act regulations, and policies of NIH, an AAALAC-accredited facility. Macaques were individually housed in 18 cu ft stainless steel cages on a 12- hour light/dark cycle in a temperature-controlled indoor facility with access to behavioral enrichment toys and were fed Purina Old World Primate Diet with rotating enrichment food. Animals were anesthetized with standard doses of Ketamine or Telazol for procedures considered to cause pain or distress to humans.

### Antibodies, cell lines, and SIV virus

The SIV Env glycoprotein-specific monoclonal antibody (mAb) clone 7D3 was produced from murine 7D3 hybridoma provided by Dr. Jim Hoxie and purified by Protein A affinity chromatography by the AIDS and Cancer Virus Program/Biological Products Core Lab of the Frederick National Laboratory for Cancer Research. SDS-PAGE analysis of the 7D3 lots demonstrated that they contained almost exclusively the antibody. Dr. Mario Roederer at the NIH/NIAID/VRC provided the rhesus ITS103.01LS mAb. Dr. Francois Villinger provided the SIV1C cell line and SIVmac239-nef-stop virus. MT4 cell line contributed by Dr. Douglas Richman (catalog #120) and SIVmac251 virus^38^ contributed by Dr. Ronald Desrosiers (catalog #253) were obtained through the NIH HIV Reagent Program, Division of AIDS, NIAID, NIH. Dr. Yoshiaki Nishimura provided the SIVmac239 virus.

The SIV1C cells, which constitutively express SIV gp120 on the cell surface and activated 24h prior with PMA (5ng/mL) and Ionomycin (500ng/mL), or MT4 cells infected with SIVmac251 or SIVmac239-nef-stop at a multiplicity of infection estimated between 0.001-0.03, for 6-15 days, with mean viability 34% (n=24 kinetics; 95% confidence interval 26%-43%) were used as SIV Env expressing cells. The mean viability of uninfected MT4 cells used as negative controls was 91% (n=15; 95% confidence interval 85%-97%).

### Conjugation and ^89^Zr radiolabeling of anti-gp120 mAbs

#### Preparation of F(ab′)_2_

We prepared ITS103.01LS-F(ab′)_2_ by pepsin digestion according to the manufacturer’s instruction (Pierce F(ab’)_2_ Preparation Kit, Thermo Fisher Scientific). Briefly, 20 mg of ITS103.01LS was digested by pepsin immobilized on agarose gel (Thermo Fisher Scientific) in 0.1M sodium acetate buffer (pH 4.4). The digestion was performed at 37°C for 1h using an optimal antibody/pepsin weight ratio between 3.5-5:1. After digestion, the reaction mixture was purified using Protein-A Sepharose affinity chromatography (Thermo Fisher Scientific), followed by dialysis in PBS at 4°C for 22h, using a dialysis membrane cassette with 20 kDa molecular weight cut-off. The purified protein was concentrated using an Amicon Ultra (MilliporeSigma) to 13 mg/mL. Protein purity was assessed by SDS-PAGE (Invitrogen) and size-exclusion HPLC (Gilson, Middleton, WI) equipped with a size-exclusion TSK gel G3000SWxL column (7.8 × 300 mm, 5 μm, TOSOH Bioscience; 0.067M sodium phosphate/0.15M sodium chloride with 0.1M KCl, pH 6.8; 1.0 ml/min) and a UV monitor.

#### Conjugation of p-SCN-Df to mAbs

The bifunctional chelating agent p- isothiocyanatobenzyl-desferrioxamine (Df-Bz-NCS) was conjugated with lysine residues of the mAbs following Vosjan et al ^39^. Briefly, for *in-vitro* assays, 200-250µg of each anti-gp120 mAb was reacted with Df-Bz-NCS at a molar ratio of 1:5 for 7D3 (or 1:3 for ITS103.01LS-F(ab′)_2_) in 0.1M sodium bicarbonate, pH 9.5, 37°C for 1 h. For *in-vivo* PET imaging, 4mg of intact 7D3 or ITS103.01LS-F(ab′)_2_ mAb was used for conjugation each time. The Df-conjugated mAbs were purified with a Zeba Spin Desalting column (7K molecular weight cutoff (MWCO), Pierce Biotechnology, Rockford, Illinois) pretreated with 0.25M sodium acetate buffer.

#### Radiolabeling

^89^Zr-Df-7D3 and ^89^Zr-Df-ITS103.01LS-F(ab′)_2_ were prepared as previously described but with minor modifications ^39^. Briefly, 37 MBq of ^89^Zr (or 370 MBq for *in-vivo*) in 1M oxalic acid was neutralized with 2M sodium bicarbonate, followed by the addition of 0.5M HEPES buffer (pH 7.1-7.3) and gentisic acid (25mg/mL, 5µL, pH 6.5). Then, 200µg of Df- conjugated mAb for *in-vitro* and 3.5 mg for *in-vivo* studies was added, and the pH was adjusted to 6.8-7.2, incubating for 1h at 30°C. The labeled product was purified by eluting with PBS 1X (pH 7.2) through a PD-10 column pretreated with 25mg BSA containing 1 µmol DTPA to block nonspecific protein binding sites. The radiochemical purity was assessed by analytical SE-HPLC (Gilson, Middleton, WI) using a column (TSK G4000PWxL (7.8×300 mm, 10μm, TOSOH Bioscience; 0.067M PBS with 0.1M KCl, pH 6.8; 1.0 mL/min) equipped with an online flow radioactivity detector (Bioscan Inc., Washington, DC). The radiolabeling yield was determined based on the distribution of ^89^Zr activities between ^89^Zr labeled mAb and unbound ^89^Zr on iTLC and/or HPLC profiles obtained before the purification. The radiochemical purity utilized in all studies was 100%.

### Binding specificity and binding affinity (K_d_) assays of radiolabeled anti-gp120 mAbs

The *in-vitro* binding specificities of ^89^Zr-7D3 and ^89^Zr-ITS103.01LS-F(ab’)_2_ to SIV Env expressing cells, cryopreserved and fresh peripheral blood mononuclear cells (PBMC), lymph node mononuclear cells (LNMC), and spleen cells of RMs were measured as previously described for anti-CD4 mAbs ^29^. Briefly, cells were washed and 1-8 million viable cells per well were suspended in 100µL (for non-specific binding) or 125µL (for total binding) of media. To measure total binding (TB), cells were incubated with 50µL of ^89^Zr-7D3 (incubation concentration 0.15nM- 6.7nM) or ^89^Zr-ITS103.01LS-F(ab’)_2_ (incubation concentration 0.5nM) for 90 min at 4°C on a rocker. To measure non-specific binding (NSB), cells were preincubated with 25µL of unlabeled 7D3 or ITS103.01LS-F(ab’)_2_ (incubation concentration 1000nM) for 20 min at 37°C in a humidified 5% CO_2_ incubator, and subsequently with 50µL of ^89^Zr-7D3 (incubation concentration 0.15nM-6.7nM) or ^89^Zr-ITS103.01LS-F(ab’)_2_ (incubation concentration 0.5nM) for 90 min at 4°C on a rocker. After the incubation, samples were microcentrifuged at 12,000g for 5 min (Eppendorf 5415C), supernatant aspirated and discarded, and the counts per minute (CPM) in the cell pellets were measured in a gamma counter (PerkinElmer 1480 Wizard or 2480 Wizard^2^). The binding specificity was expressed as the CPM ratio of TB to NSB. In one assay where the NSB wells were missing due to an insufficient amount of unlabeled mAb, the TB of uninfected MT4 cells was substituted for NSB to estimate the binding specificity of the radioligand to SIV Env expressing cells.

The media used is as follows: RPMI-1640 without L-Glutamine, 10% Heat Inactivated FBS, and supplemented with Penicillin-Streptomycin for SIV1C; RPMI-1640 without L-Glutamine, 10% Heat Inactivated FBS, 1% L-Glutamine, 1% Hepes Buffer, and supplemented with Gentamicin for MT4; PBS (phosphate buffered saline) pH 7.4 for PBMC, LNMC, and spleen cells.

A secondary binding specificity assessment was performed by measuring the interaction between ^89^Zr-7D3 or ^89^Zr-ITS103.01LS-F(ab’)_2_ and SIV gp120 (mac239) protein (eEnzyme LLC). Immuno-breakable well strips (Thermo Scientific) were coated with 150µl of SIV gp120 (4.0 µg/ml in phosphate-buffered saline (PBS), pH 7.2), incubated overnight at 4°C, and then washed with 0.05% Tween-20 in PBS (Washing buffer, Sigma Aldrich). The wells were then preincubated with either 200µl of BSA blocking buffer (Thermo Scientific; 5% bovine serum albumin in PBS) for total binding (TB) or 200µl of unlabeled 7D3 or ITS103.01LS-F(ab’)_2_ (1000nM in BSA blocking buffer) for non-specific binding (NSB) for 1h at 37°C. The wells were then washed and incubated at various concentrations [0.05nM-50nM] of 150µl of ^89^Zr-7D3 or ^89^Zr-ITS103.01LS- F(ab’)_2_ for an additional 1h at 37°C. After the incubation, the wells were washed, the dried breakable wells were detached individually, and the radioactivity (CPM) in each well was measured using a gamma counter (PerkinElmer 2480 Wizard^2^). The wells were washed thrice with 200µl of the Washing buffer during each washing step. The binding specificity was expressed as the CPM ratio of TB to NSB. Specific binding was calculated by subtracting the NSB CPM from the TB CPM. The binding affinity, represented by the equilibrium disassociation constant K_d_ and equal to the free ligand concentration at 50% of maximal binding was estimated using GraphPad Prism 9.3.1 (La Jolla, CA) by fitting the one-site specific binding curve with a weight of 1/y^2^.

### Pre-existing immune response against anti-gp120 mAbs

The presence of any pre-existing immune response against 7D3 or ITS103.01LS was tested with radio-HPLC before administration of the radiolabeled mAb for PET/CT imaging as previously described ^40^. Briefly, monkey plasma was incubated with ^89^Zr-7D3 or ^89^Zr-ITS103.01LS-F(ab’)_2_ (mAb concentration 1.5nM-5nM) for 30 min at 37°C in a humidified 5% CO_2_ incubator, and a 50µl aliquot of the incubated mixture was run through size-exclusion HPLC.

### Endogenous proteins competing for the anti-gp120 mAb binding site and immunogenicity of murine 7D3 or rhesus ITS103.01LS-F(ab’)_2_

The presence of endogenous proteins in the plasma that compete for 7D3 or ITS103.01LS-F(ab’)_2_ binding site and the development of immunogenic response post-exposure against the 7D3 or ITS103.01LS-F(ab’)_2_ mAb were detected with a cell binding assay coupled with radio-HPLC as previously described ^40^. Briefly, monkey plasma was incubated with ^89^Zr-7D3 or ^89^Zr- ITS103.01LS-F(ab’)_2_ (mAb concentration 1.5nM-5nM) for 30 min at 37°C in a humidified 5% CO_2_ incubator. After incubation, a 50µl aliquot of the incubated mixture was run through size-exclusion HPLC, and a 20µl aliquot of the incubated mixture was added to 1-2 million viable gp120 expressing cells in 180µL. After 90 min incubation (mAb concentration 0.15nM-0.5nM) on a rocker at 4°C, the total incubated radioactivity CPM was measured in a gamma counter (PerkinElmer 1480 Wizard or 2480 Wizard^2^). The cell mixture was then microcentrifuged at 12,000g for 5 min (Eppendorf 5415C), supernatant aspirated and discarded, the CPM in the cell pellet measured, and the percent of total incubated radioactivity bound to gp120 expressing cells was determined.

### Stability and binding ability of radiolabeled anti-gp120 mAbs in *in-vivo* plasma

For the *in-vivo* stability test, 50µl of the plasma obtained ∼13h and ∼40h post ^89^Zr-7D3 injection or ∼9h post ^89^Zr-ITS103.01LS-F(ab’)_2_ injection was run through size-exclusion HPLC. For the *in-vivo* plasma binding ability test, a binding assay with radioactive plasma obtained at 1h, 2h, 4h, 6h, or ∼13h post ^89^Zr-7D3 injection or ∼9h post ^89^Zr-ITS103.01LS-F(ab’)_2_ injection was performed with SIV gp120 expressing cells or SIV gp120 protein coated on wells as described above in the Binding specificity section. Briefly, 1 million viable SIV gp120 expressing cells were incubated with 50µl of *in-vivo* radioactive plasma following the procedure described above to assess total binding and non-specific binding. Separately, SIV gp120 coated wells were first blocked for non-specific binding with BSA blocking buffer or with unlabeled 7D3 or ITS103.01LS-F(ab’)_2_ (1µM in BSA blocking buffer) for 1h at 37°C, and following washing and incubating with 150µl of *in-vivo* radioactive plasma for an additional 1h at 37°C, following the method described above for secondary binding specificity test. The binding specificity was expressed as the CPM ratio of TB to NSB.

### *In-vitro* autoradiography

20µm thick slices of axillary and inguinal lymph nodes, spleen, colon, and jejunum tissues were sectioned with a cryostat (Leica Biosystems CM1900 UV), mounted on silanized slides, dried at room temperature for 1-2h, and stored at -80°C when not immediately used. The fresh slides were dried for 1-2h whereas slides stored at -80°C were air-dried overnight at room temperature before use. The dried slides were pre-incubated in the buffer for 15 min at room temperature. To measure total binding (TB), slides were incubated with 1nM of ^89^Zr-7D3 or ^89^Zr-ITS103.01LS-F(ab’)_2_ for 90 min at 4°C. To determine non-specific binding (NSB), the slides were preincubated with 100nM of unlabeled 7D3 or ITS103.01LS-F(ab’)_2_ for 20 min at room temperature and subsequently incubated with 1nM of ^89^Zr-7D3 or ^89^Zr-ITS103.01LS-F(ab’)_2_ for 90 min at 4°C. After incubation, slides were rinsed twice (5 min per rinse) in cold buffer and air-dried for 1h. All incubation and rinsing steps were performed with 50mM Tris-HCL buffer (pH 7.4). The dried slides were exposed to a storage phosphor screen along with ^89^Zr-7D3 or ^89^Zr-ITS103.01LS-F(ab’)_2_ standards. After 20-24h of exposure, the phosphor screen was read using a phosphorimager (GE Healthcare Typhoon FLA 7000) at 25µm pixel resolution. Regions of interest (ROIs) were drawn manually on the images and the average intensity was extracted from each tissue using image analysis software (GE Healthcare ImageQuant TL). The binding specificity is calculated as the average intensity ratio of TB to NSB.

### PET/CT imaging and data analysis

Animals were initially anesthetized with a restraint dose of ketamine (10 mg/kg, intramuscular) and after shaving the skin and prepping the insertion site, a 22-gauge catheter was inserted in the saphenous vein of the leg for bolus injection of the radiotracer or continuous administration of anesthetics (propofol 0.2 mg/kg/min infusion) during the imaging procedure. The macaque’s arms and legs were restrained and positioned in supine orientation for imaging. Anesthetized primates were monitored with a pulse oximeter and thermometer, and the body temperature during imaging was maintained with Bair Hugger patient warming system.

*In-vivo* PET imaging was performed with MultiScan LFER 150 PET/CT camera (Mediso Medical Imaging Systems, Budapest, Hungary) designed for nonhuman primates. Whole-body CT acquisitions were performed using the following parameters: Semi-circular multi field-of-view (FOV), 360 projections, 80kVp, 710µA, 90ms exposure time per projection, and 1:4 binning. Immediately following the CT scan, animals underwent a whole-body static PET scan from the top of the head to mid-thighs at one or multiple time points post radioligand injection (Supplementary Table 1 and 2). PET static scans were acquired at 10 min per FOV, and 6-bed positions were scanned with a 35% overlap between the FOVs with 1-9 coincidence mode and a 5 ns coincidence time window. Raw CT data was reconstructed with scatter correction. Raw PET data was reconstructed using the following parameters: 400-600 keV energy window, 1-9 coincidence mode, Tera-Tomo 3D reconstruction with median and spike filter on, voxel size 1 mm, and 8 iterations and 9 subsets. Reconstructed PET images were corrected for attenuation (using CT material map segmentation), radioactive decay, uniformity, random coincidences, scattering of radiation, and the decay reference was set to radioligand administration time. A Gaussian post-processing filter with kernel size = 3 and sigma = 0.8 was applied to smooth the reconstructed PET image. Final CT and PET images were saved in DICOM format.

PET images were analyzed using MIM software (Cleveland, USA), independently and blindly by three operators (authors: SS, HJ, and JAC). Volumes of interest (VOI) on the co-registered PET and CT image were manually drawn for cervical, axillary, and inguinal lymph nodes (LNs), spleen, heart (blood pool), liver, left kidney, bone marrow, muscle, and gut, and the amount of radioligand uptake was quantified as maximum and/or mean standardized uptake value (SUV) using the formula: SUV = (*c/d)*w*, where *c* is decay-corrected tissue concentration (µCi/mL), *d* is the injected dose (µCi), and *w*, the body weight (g). Bone marrow uptake was evaluated at three locations: L4 lumbar vertebrae, entire thoracic and lumbar vertebrae, and the proximal humerus. Muscle uptake was measured by drawing VOI on the back muscles adjacent to the spine at the axial level of the heart. Care was taken to avoid any abdominal LNs in the gut VOI. Images and the quantitative measures were not corrected for the partial volume effect.

Whole blood and plasma radioactivity CPM were measured by drawing blood and counting 50µL or 100µL aliquots in a gamma counter. CPM/mL was then converted to SUV correcting for radioactive decay and adjusting for counting efficiency, body weight, and injected radioactivity. Whole blood and plasma SUV were calculated at each imaging timepoint and additionally at ∼13h post-injection (for ^89^Zr-7D3) or ∼9h and ∼24h post-injection (for ^89^Zr-ITS103.01LS-F(ab’)_2_).

### SUV normalization

To control for differences in the clearance of the radioligand from the intravascular compartment between the animals, blood-adjusted tissue uptake (rSUVmax and rSUVmean) was calculated by normalizing the tissue uptake on blood SUV obtained from heart blood pool VOI ie SUVmax or mean of tissue/SUVmean of blood pool. The heart blood pool SUVmean extracted from the 48 PET images correlated strongly with whole-blood and plasma SUV calculated from counting aliquots in a gamma counter (ρ ≥0.92, P <0.0001). Hence, the cardiac blood pool SUVmean from the PET image was used for normalization.

### Ex-vivo analysis

One chronically SIV-infected RM and one uninfected RM pair at ∼44h post ^89^Zr-7D3 injection, another pair on Day 7 post ^89^Zr-7D3 injection, and another pair on Day 5 post ^89^Zr-ITS103.01LS- F(ab’)_2_ injection were euthanized and individual LNs (axillary, inguinal, submandibular, mesenteric, and retroperitoneal), and small aliquots of the spleen and gut sections (duodenum, jejunum, ileum, and colon) after removal of content were harvested, weighed, radioactivity measured in a gamma counter (PerkinElmer 1480 Wizard or 2480 Wizard^2^), and decay-corrected tissue radioactivity concentration (µCi/g) and SUVmean were calculated.

### Lymphocyte immunophenotyping, plasma viral load, and cell-associated viral load

Fresh blood samples collected in EDTA tubes were stained for CD3 and CD4, and analyzed by flow cytometry as previously described ^41^. Plasma SIV-RNA was measured using a gag-targeted quantitative real-time/digital reverse transcription-polymerase chain reaction (RT-PCR) assay with a minimum detection threshold of 3 or 5 copies/mL as previously described ^42^. Cell-associated viral load from six independent aliquots (each one run in duplicates) of SIVmac251 or SIVmac239-nef-stop viral kinetics in MT4 cells was measured and expressed as SIV-RNA copies/million cells, as previously described ^43^.

### Statistical analysis

Unpaired samples were compared with the non-parametric Wilcoxon rank-sum test and the relationship between variables in the cross-sectional analysis was assessed with Spearman rank correlation using Winstat software (Cambridge, MA). A P-value <0.05 was considered statistically significant.

## List of supplementary materials

Supplementary figures 1-6 and Supplementary tables 1-7.

## Acknowledgments

We thank the animal care staff and technicians at the NIH Animal Center (Dickerson, Maryland, USA) for their care and handling of the animals, Dr. Jeffrey Lifson, Julian Bess, and Anitha Vijayagopalan for providing the 7D3 mAb, Dr. Mario Roederer for providing the ITS103.01LS mAb, Dr. Ken Matsui, Dr. Hiromi Imamichi and Mindy Smith for technical assistance, and Dr. Marisa St. Claire for veterinary support and technical assistance throughout the study. We thank Dr. Cliff Lane for his ongoing support and useful discussions on the study. We thank Dr. Paolo Lusso for the initial discussions on the study design.

## Author contributions

S.S., I.K., J.A.C., and M.D.M. conceived the study and designed the research.

S.S. and M.D.M. wrote the manuscript with assistance from I.K. and J.A.C. S.S., H.J., and M.D.M. made the figures and tables.

S.S. performed statistical analyses.

S.S., I.K., H.J., P.D., H.B., V.D., Y.B., V.N., and B.L. performed research.

M.D.M. provided oversight of the study.

All authors analyzed and interpreted the data, and critically reviewed the manuscript.

