## Supplemental data for "Whole-body PET imaging of SIV using anti-Env probes fails to reveal regions of specific uptake in rhesus macaques"

**Running title:** immunoPET imaging of SIV
**Keywords:** immunoPET, rhesus macaques, endogenous antibodies, 7D3, ITS103, reproducibility

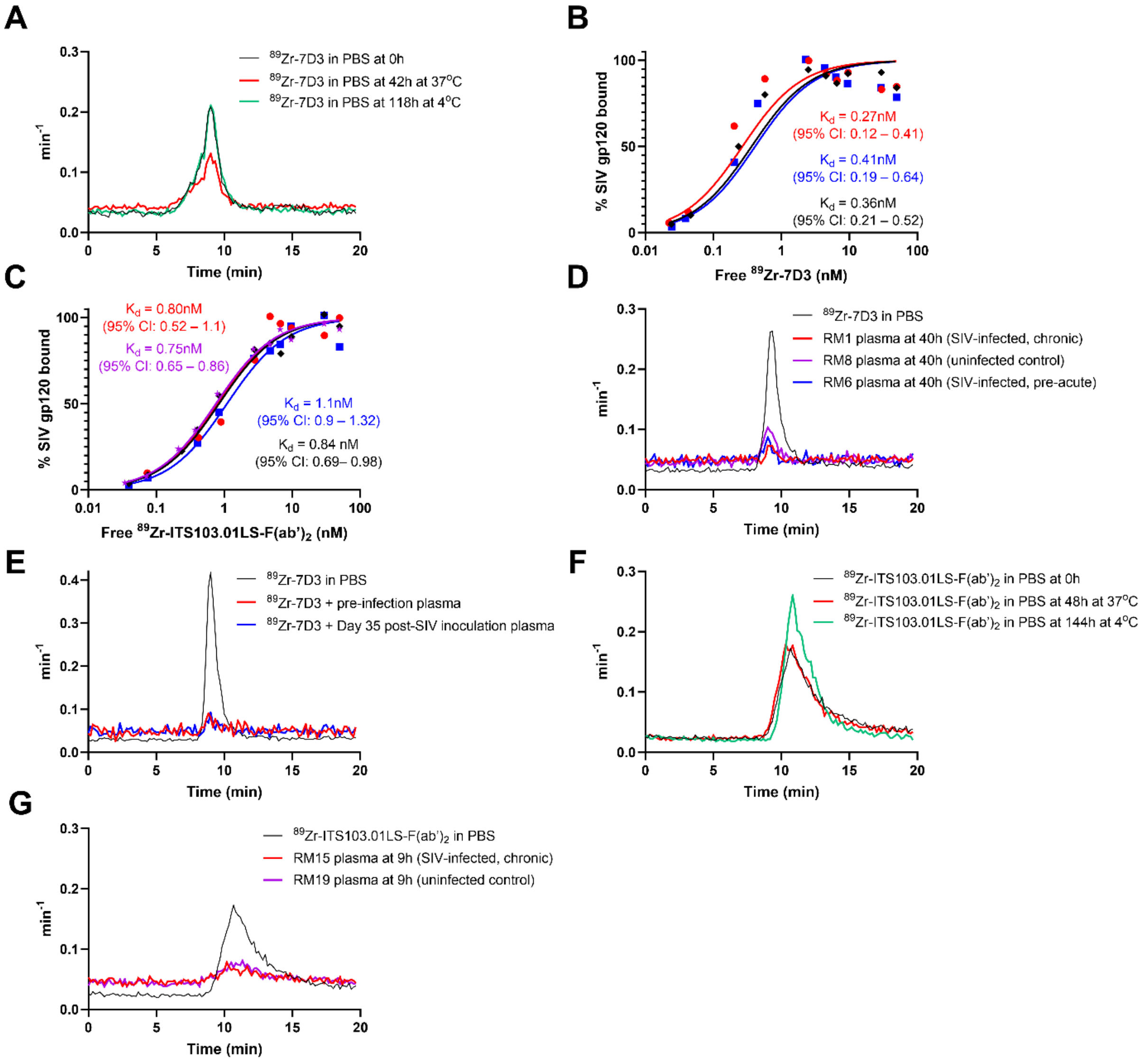

**Supplementary Fig. 1** Stability of $^{89}$Zr radiolabeled 7D3 in PBS (black) assessed through radio-HPLC at 42h (red, radioligand stored at 37°C) and at 118h (green, radioligand stored at 4°C) post-radiolabeling (**A**). The semi-logarithmic plots of saturation binding curves of $^{89}$Zr-7D3 (**B**) and $^{89}$Zr-ITS103.01LS-F(ab')$_2$ (**C**). Representative radio-HPLC profiles of the plasmas obtained from animals at 40h post $^{89}$Zr-7D3 injection confirmed near 100% stability of $^{89}$Zr-7D3 *in-vivo* **(D)**. Radio-HPLC of the incubated $^{89}$Zr-7D3 and plasma at pre-infection and Day 35 post-SIVmac239-nef-stop inoculation in a representative macaque **(E)**. The stability of $^{89}$Zr radiolabeled ITS103.01LS-F(ab')$_2$ in PBS (black) was assessed through radio-HPLC at 48h (red, radioligand stored at 37°C) and at 144h (green, radioligand stored at 4°C) post-radiolabeling (**F**). Representative radio-HPLC profiles of the plasmas obtained from animals at 9h post $^{89}$Zr-ITS103.01LS-F(ab')$_2$ injection confirmed near 100% stability of $^{89}$Zr-ITS103.01LS-F(ab')$_2$ *in-vivo* **(G)**. All radiochromatograms were transformed into probability density curves by normalizing the area under the curve.

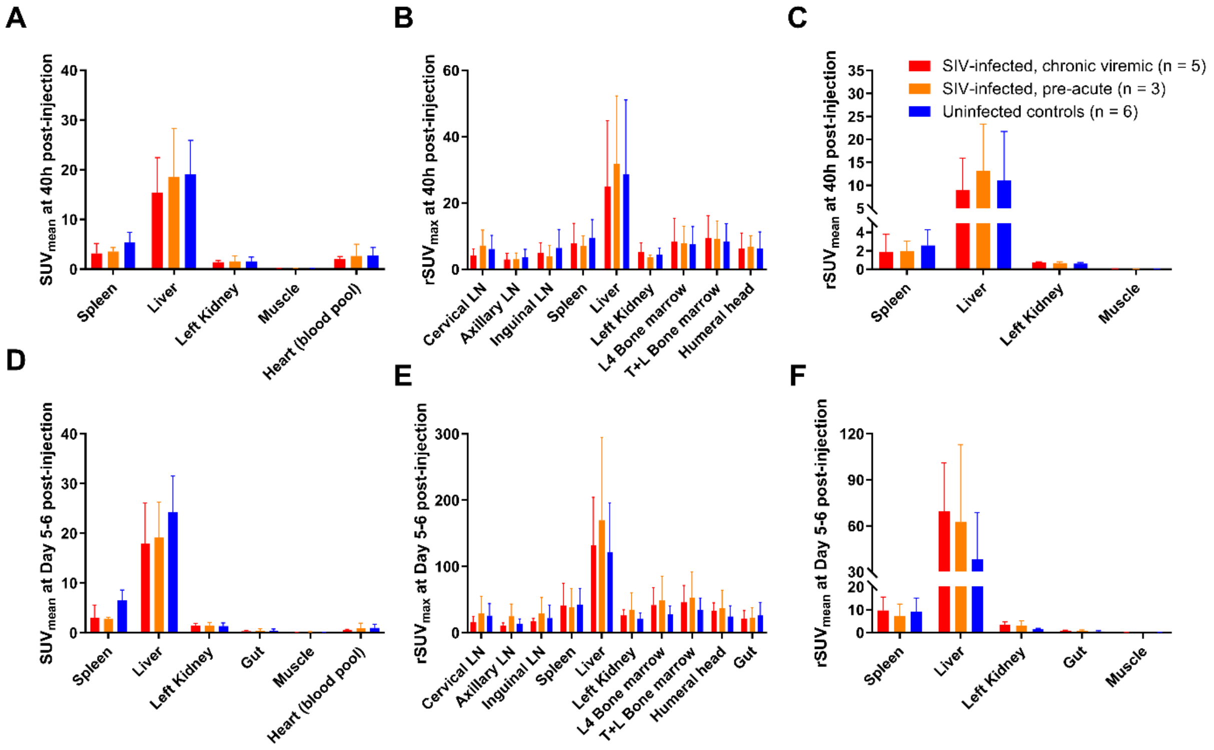

**Supplementary Fig. 2** Comparison of mean ($SUV_{mean}$), blood-adjusted maximum ($rSUV_{max}$), and blood-adjusted mean ($rSUV_{mean}$) standardized uptake value (SUV) in tissues among the chronically SIV-infected viremic (red), pre-acutely SIV-infected (orange), and uninfected controls (blue) at 40h (**A-C**) and Day 5-6 post $^{89}Zr$-7D3 injection (**D-F**). Plots are mean values and error bars are standard deviation.

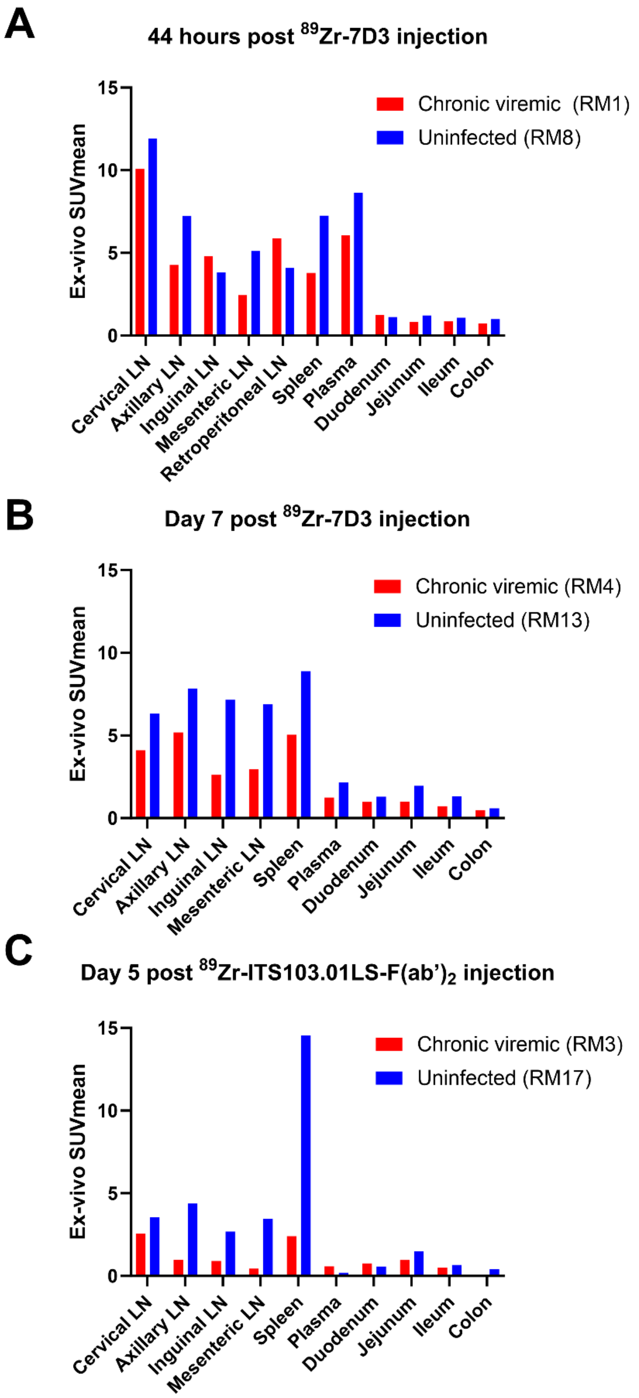

**Supplementary Fig. 3** SUV in necropsy tissues obtained from one chronically SIV-infected RM (red) and one uninfected control (blue) at ~44h (**A**) and Day 7 post $^{89}$Zr-7D3 injection (**B**), and Day 5 post $^{89}$Zr-ITS103.01LS-F(ab')$_2$ injection (**C**).

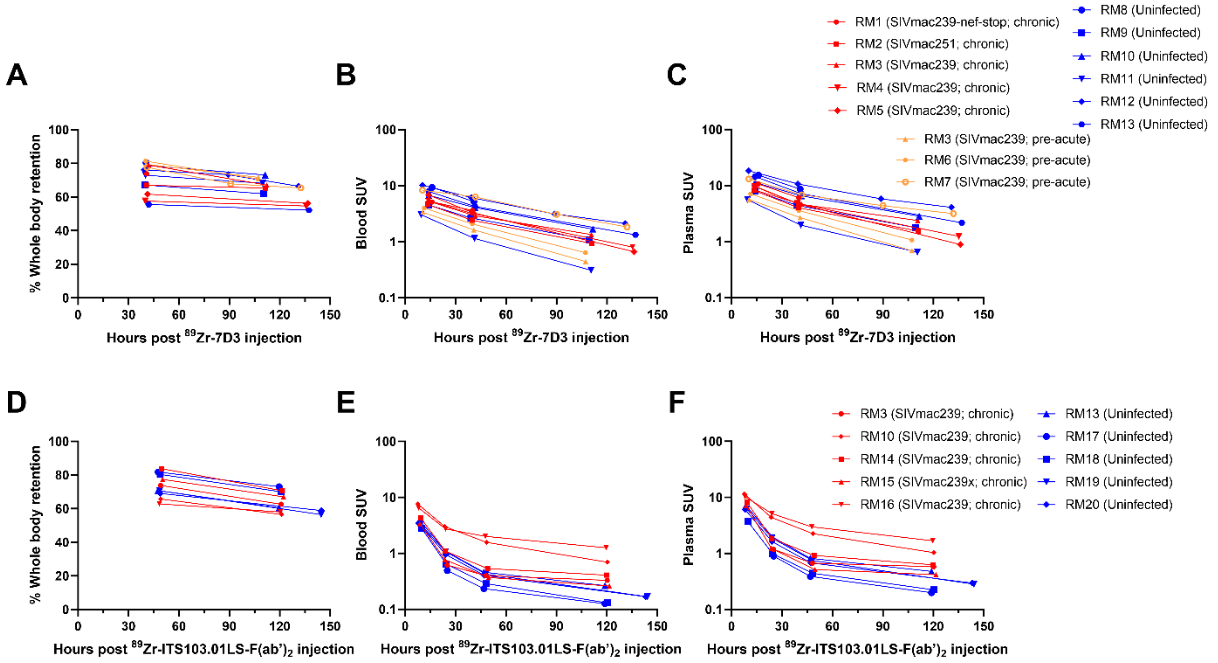

**Supplementary Fig. 4** Whole-body (head to mid-thighs) clearance and standardized uptake values (SUV) in the peripheral blood and plasma of $^{89}$Zr-7D3 (**A, B, C**) and $^{89}$Zr-ITS103.01LS-F(ab')$_2$ (**D, E, F**).

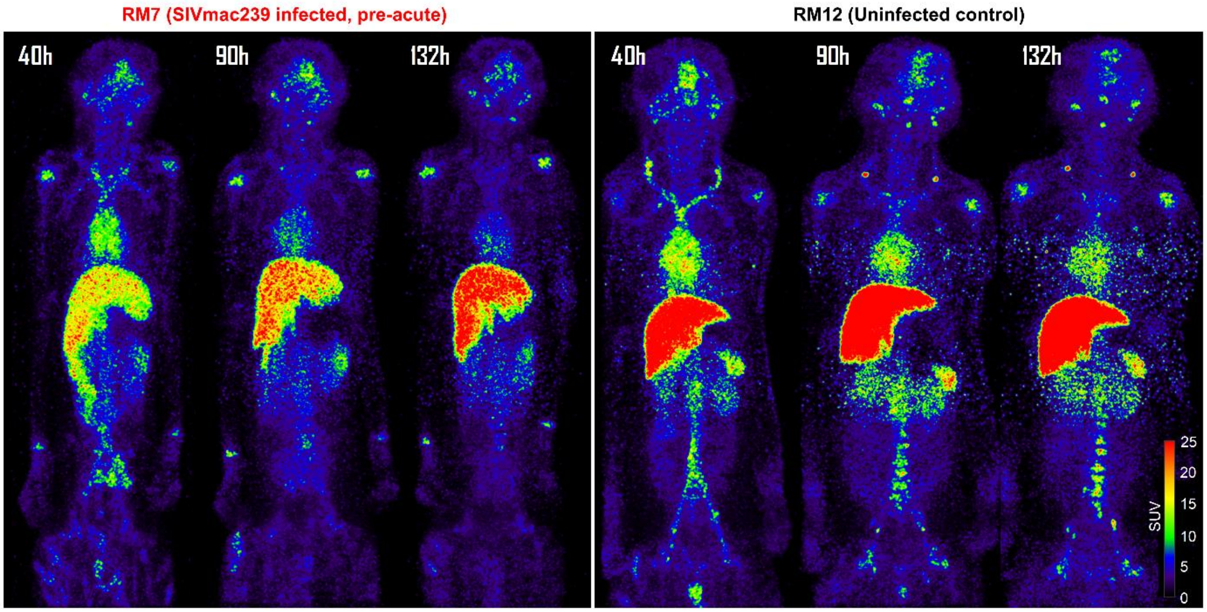

**Supplementary Fig. 5** Maximum intensity projection PET images of a representative pre-acutely SIV-infected rhesus macaque (RM7) and a representative uninfected control (RM12) following administration of ~1mg mass of $^{89}$Zr-7D3 (~0.4mCi of $^{89}$Zr) and scanned at 40h, 90h, and Day 6 post-injection **(A)**. The SIV-infected animal was inoculated intravenously with 1000 TCID50 SIVmac239, administered $^{89}$Zr-7D3 on Day 8 of SIV-infection, and PET/CT scanned on Day 10 (40h), Day 12 (90h), and Day 14 (132h) of infection. Tissue uptakes were displayed on the RAINBOW color scale where the red color indicates a high standardized uptake value (SUV). Both visual and semi-quantitative SUV analysis showed similar uptake in the SIV-infected RM compared to the uninfected control and the gut uptake observed at 40h had cleared at later timepoints.

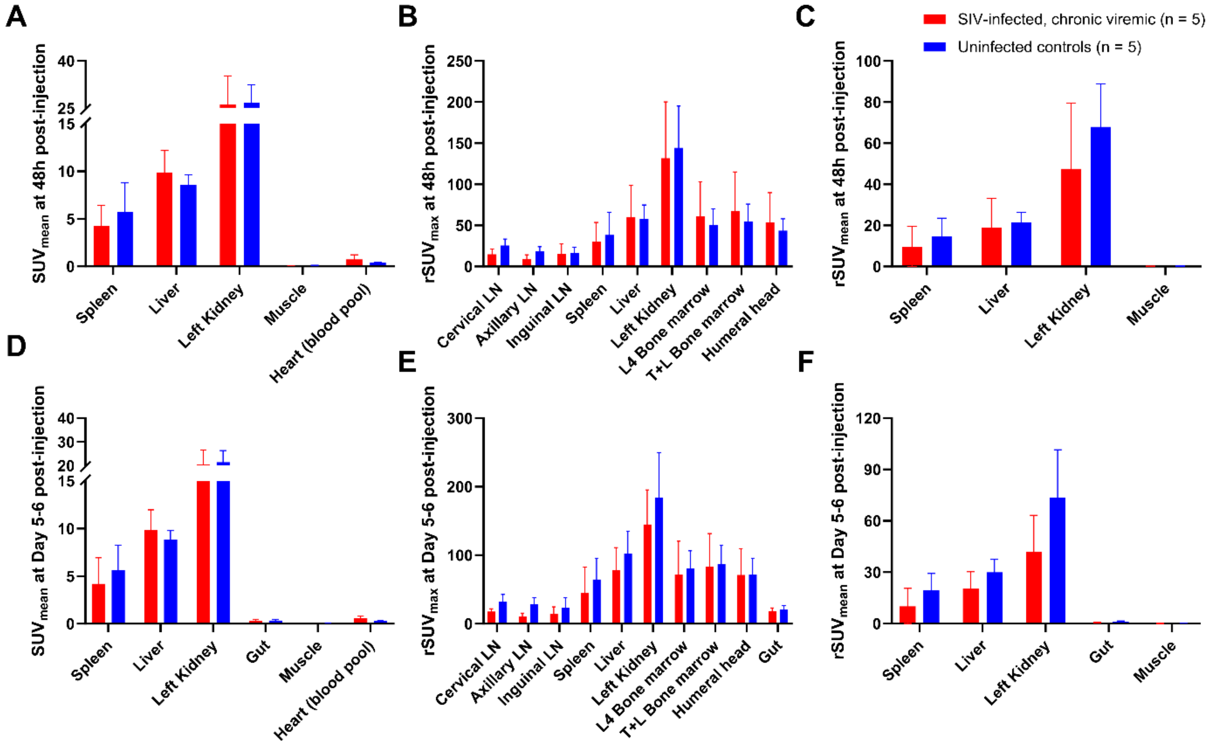

**Supplementary Fig. 6** Comparison of mean ($SUV_{mean}$), blood-adjusted maximum ($rSUV_{max}$), and blood-adjusted mean ($rSUV_{mean}$) standardized uptake value (SUV) in tissues among the chronically SIV-infected viremic (red) and uninfected controls (blue) at 48h (**A-C**) and Day 5-6 post $^{89}$Zr-ITS103.01LS-F(ab')$_2$ injection (**D-F**). Plots are mean values and error bars are standard deviation.

**Supplementary Table 1.** Characteristics of Indian Rhesus Macaques used for *in-vivo* PET/CT imaging with $^{89}$Zr-7D3.

| Stage of infection on $^{89}$Zr-7D3 injection | Macaque designation | SIV strain/infection route/$TCID_{50}$ | PET/CT imaging post injection (hours) | Gender | Age (Y) | Body Weight (kg) | Plasma viral load (copies/mL)* | Peripheral blood CD4 (cells/µL) | 7D3 mass (µg) | Inj. $^{89}$Zr Activity (mCi) |
|---|---|---|---|---|---|---|---|---|---|---|
| Chronic, 24 months off ART | RM1 | SIVmac239-nef-stop/IV/200 | 41 | Male | 8.9 | 10.05 | $8.9 \times 10^5$ | 2 | 636 | 2.07 |
| Chronic, 7 months, ART naïve | RM2 | SIVmac251/IV/300 | 41, 112 | Male | 7.7 | 13.2 | $1.1 \times 10^5$ | 153 | 942 | 1.94 |
| Chronic, 18 months, ART naïve | RM3 | SIVmac239/IV/300 | 42, 112 | Male | 6.5 | 10.45 | $1.3 \times 10^6$ | 147 | 929 | 0.77 |
| Chronic, 58 months, ART naïve | RM4 | SIVmac239/IV/500 | 40, 136 | Male | 8.0 | 7.85 | $1.2 \times 10^4$ | 21 | 1008 | 2.23 |
| Chronic, 58 months, ART naïve | RM5 | SIVmac239/IV/500 | 41, 137 | Female | 9.0 | 5.85 | $1.6 \times 10^6$ | 284 | 1000 | 2.14 |
| Pre-acute, Day 8 | RM3 | SIVmac239/IV/300 | 41, 108 | Male | 5.0 | 8.6 | $2.6 \times 10^3$ - $2.5 \times 10^6$ | # | 1007 | 2.02 |
| Pre-acute, Day 8 | RM6 | SIVmac239/IV/300 | 40, 107 | Male | 7.3 | 9.45 | $3.6 \times 10^3$ - $1.2 \times 10^6$ | # | 970 | 1.98 |
| Pre-acute, Day 8 | RM7 | SIVmac239/IV/1000 | 41, 91, 132 | Male | 5.5 | 6.9 | $1.2 \times 10^7$ - $4.8 \times 10^6$ | # | 1040 | 0.45 |
| Uninfected control | RM8 | NA | 40 | Male | 5.9 | 7.15 | NA | 438 | 644 | 2.12 |
| Uninfected control | RM9 | NA | 40, 110 | Male | 7.4 | 12.25 | NA | 606 | 936 | 1.93 |
| Uninfected control | RM10 | NA | 41, 111 | Male | 6.4 | 11.8 | NA | 1328 | 962 | 0.8 |
| Uninfected control | RM11 | NA | 41, 114 | Male | 5.0 | 7.95 | NA | 871 | 982 | 1.94 |
| Uninfected control | RM12 | NA | 40, 90, 131 | Male | 7.4 | 10 | NA | 774 | 952 | 0.41 |
| Uninfected control | RM13 | NA | 42, 137 | Male | 7.8 | 9.6 | NA | 872 | 933 | 1.99 |

$TCID_{50}$, median tissue culture infectious dose; IV, intravenous; ART, antiretroviral therapy; NA, not applicable

* Plasma viral load ranging from Day 7 to Day 14 post-SIV inoculation was reported for pre-acute RMs.

### Cell counts of pre-acute RMs between Day 7 and Day 14 post-SIV inoculation are not available.

**Supplementary Table 2.** Characteristics of Indian Rhesus Macaques used for *in-vivo* PET/CT imaging with $^{89}$Zr-ITS103.01LS-F(ab')$_2$.

| Stage of infection on $^{89}$Zr-ITS103.01LS-F(ab')$_2$ injection | Macaque designation | SIV strain/infection route/TCID$_{50}$ | PET/CT imaging post injection (hours) | Gender | Age (Y) | Body Weight (kg) | Plasma viral load (copies/mL) | Peripheral blood CD4 (cells/µL) | ITS103.01LS-F(ab')$_2$ mass (µg) | Inj. $^{89}$Zr Activity (mCi) |
|---|---|---|---|---|---|---|---|---|---|---|
| Chronic, 20 months, ART naïve | RM3 | SIVmac239/IV/300 | 49, 121 | Male | 6.6 | 9.95 | $4.4 \times 10^6$ | 137 | 977 | 1.67 |
| Chronic, 7 weeks, ART naïve | RM10 | SIVmac239/IV/1000 | 49, 121 | Male | 6.8 | 9.95 | $2.6 \times 10^7$ | 948 | 963 | 1.76 |
| Chronic, 12 months, ART naïve | RM14 | SIVmac239/IV/3000 | 50, 122 | Female | 14.4 | 7.96 | $1.0 \times 10^4$ | 250 | 966 | 1.58 |
| Chronic, 17 months, ART naïve | RM15 | SIVmac239x/Rectal/4 | 50, 122 | Male | 6.8 | 7.45 | $3.0 \times 10^6$ | 315 | 1041 | 1.70 |
| Chronic, 7 weeks, ART naïve | RM16 | SIVmac239/IV/1000 | 48, 120 | Male | 7.2 | 8.95 | $2.5 \times 10^7$ | 377 | 907 | 1.73 |
| Uninfected control | RM13 | NA | 48, 120 | Male | 7.0 | 9.25 | NA | 602 | 942 | 1.80 |
| Uninfected control | RM17 | NA | 48, 120 | Male | 2.7 | 4.20 | NA | 2670 | 972 | 1.66 |
| Uninfected control | RM18 | NA | 49, 121 | Male | 2.8 | 4.65 | NA | 2733 | 1047 | 1.72 |
| Uninfected control | RM19 | NA | 48, 145 | Male | 4.8 | 5.25 | NA | 3353 | 998 | 2.43 |
| Uninfected control | RM20 | NA | 49, 145 | Male | 3.8 | 5.4 | NA | 1981 | 983 | 2.37 |

TCID$_{50}$, median tissue culture infectious dose; IV, intravenous; ART, antiretroviral therapy; NA, not applicable

**Supplementary Table 3.** Rhesus macaques used for *in-vitro* 7D3 primary cell binding studies.

| | Macaque designation | Fresh/Cryo | Plasma viral load (copies/mL) | Weeks of untreated infection | ART status | SIV strain | Binding specificity |
|---|---|---|---|---|---|---|---|
| PBMC | RM14 | Fresh | $6.8 \times 10^4$ | 98 | ART naïve | SIVmac239 | 1.74 |
| | RM21 | Cryo | $3.1 \times 10^4$, $5.3 \times 10^4$ | 1, 5 | ART naïve | SIVmac239-nef-stop | 0.67, 1.15 |
| | RM22 | Cryo | $1.5 \times 10^4$ | 1 | ART naïve | SIVmac239-nef-stop | 0.6 |
| | RM23 | Cryo | $2.7 \times 10^6$ | 1.3, 2.1 | ART naïve | SIVmac239 | 1.03, 0.7 |
| | RM24 | Cryo | $1.4 \times 10^6$ | 2.1 | ART naïve | SIVmac239 | 0.67 |
| | RM25 | Cryo | $1.7 \times 10^4$ | 1 | ART naïve | SIVmac239-nef-stop | 1.29 |
| | RM26 | Fresh | $1.8 \times 10^2$ | 317 | off ART | SIVmac239-nef-stop | 1.09 |
| | RM27 | Cryo | $3.5 \times 10^4$ | 1 | ART naïve | SIVmac239-nef-stop | 0.95 |
| | RM28 | Cryo | $2.0 \times 10^5$ | 5 | ART naïve | SIVmac239-nef-stop | 0.92 |
| | RM29 | Cryo | $1.2 \times 10^4$ | 1 | ART naïve | SIVmac239-nef-stop | 1.13 |
| | RM30 | Cryo | $1.3 \times 10^4$ | 1 | ART naïve | SIVmac239-nef-stop | 0.8 |
| | RM31 | Cryo | $3.1 \times 10^4$, $1.3 \times 10^6$ | 1, 5 | ART naïve | SIVmac239-nef-stop | 0.85, 1.4 |
| | RM32 | Cryo | $3.2 \times 10^3$ | 1 | ART naïve | SIVmac239-nef-stop | 1.43 |
| | RM33 | Fresh | - | Uninfected control | - | - | 2.27 |
| | RM34 | Fresh | - | Uninfected control | - | - | 1.26 |
| Inguinal LNMC | RM15 | Cryo | $3.9 \times 10^7$ | 105 | ART naïve | SIVmac239x | 0.52 |
| | RM29 | Cryo | $2.0 \times 10^6$ | 73 | off ART | SIVmac239-nef-stop | 1.09 |
| | RM35 | Cryo | - | Uninfected control | - | - | 0.68 |
| Mesenteric LNMC | RM10 | Cryo | $2.7 \times 10^8$ | 12 | ART naïve | SIVmac239 | 1.79 |
| | RM15 | Cryo | $3.9 \times 10^7$ | 105 | ART naïve | SIVmac239x | 0.76 |
| | RM36 | Cryo | $1.4 \times 10^8$ | 10 | ART naïve | SIVmac239 | 1.00 |
| Spleen cells | RM10 | Cryo | $2.7 \times 10^8$ | 12 | ART naïve | SIVmac239 | 1.02 |
| | RM15 | Cryo | $3.9 \times 10^7$ | 105 | ART naïve | SIVmac239x | 1.16 |
| | RM36 | Cryo | $1.4 \times 10^8$ | 10 | ART naïve | SIVmac239 | 0.94 |

PBMC, peripheral blood mononuclear cells; LNMC, lymph node mononuclear cells; ART, antiretroviral therapy

**Supplementary Table 4.** Rhesus macaques used for 7D3 autoradiography studies.

| Tissue | Macaque designation | Plasma viral load (copies/mL) | Weeks of untreated infection | ART status | SIV strain | Binding specificity |
|---|---|---|---|---|---|---|
| **Axillary lymph node** | RM1 | $1.2 \times 10^7$ | 40 | off ART | SIVmac239-nef-stop | 1.1 |
| | RM6 | $9.9 \times 10^4$ | 77 | ART naïve | SIVmac239 | 1.1 |
| | RM11 | $1.2 \times 10^7$ | 32 | ART naïve | SIVmac239 | 1.0 |
| | RM37 | $1.4 \times 10^5$ | 240 | off ART | SIVmac239-nef-stop | 0.8 |
| | RM46 | $7.4 \times 10^5$ | 49 | off ART | SIVmac251 | 0.9 |
| | RM17 | - | Uninfected control | - | - | 1.1 |
| **Colon** | RM6 | $9.9 \times 10^4$ | 77 | ART naïve | SIVmac239 | 1.1 |
| | RM11 | $1.2 \times 10^7$ | 32 | ART naïve | SIVmac239 | 0.8 |
| | RM16 | $1.4 \times 10^8$ | 10 | ART naïve | SIVmac239 | 1.1 |
| | RM37 | $1.4 \times 10^5$ | 240 | off ART | SIVmac239-nef-stop | 0.9 |
| **Inguinal lymph node** | RM6 | $9.9 \times 10^4$ | 77 | ART naïve | SIVmac239 | 1.2 |
| | RM38 | $4.1 \times 10^4$ | 230 | off ART | SIVmac239-nef-stop | 1.0 |
| **Jejunum** | RM3 | $4.4 \times 10^6$ | 86 | ART naïve | SIVmac239 | 1.0 |
| **Spleen** | RM6 | $9.9 \times 10^4$ | 77 | ART naïve | SIVmac239 | 1.4 |
| | RM11 | $1.2 \times 10^7$ | 32 | ART naïve | SIVmac239 | 1.6 |
| | RM25 | $5.2 \times 10^6$ | 45 | off ART | SIVmac239-nef-stop | 1.1 |
| | RM37 | $1.4 \times 10^5$ | 240 | off ART | SIVmac239-nef-stop | 0.5 |
| | RM38 | $4.1 \times 10^4$ | 230 | off ART | SIVmac239-nef-stop | 0.7 |
| | RM46 | $7.4 \times 10^5$ | 49 | off ART | SIVmac251 | 0.8 |
| | RM17 | - | Uninfected control | - | - | 0.7 |
| | RM47 | - | Uninfected control | - | - | 0.6 |

ART, antiretroviral therapy

**Supplementary Table 5.** SIV-infected rhesus macaques used to test the development of endogenous proteins competing for the 7D3 binding site.

| Macaque designation | SIV strain/infection route/$TCID_{50}$ | Time point | Plasma viral load (copies/mL) |
|---|---|---|---|
| RM1 | SIVmac239-nef-stop/IV/200 | 105 weeks off ART | $8.9 \times 10^5$ |
| RM3 | SIVmac239/IV/300 | 77 weeks off ART | $1.3 \times 10^6$ |
| RM4 | SIVmac239/IV/500 | 253 weeks ART naïve | $1.2 \times 10^4$ |
| RM5 | SIVmac239/IV/500 | 253 weeks ART naïve | $1.6 \times 10^6$ |
| RM25 | SIVmac239-nef-stop/IV/200 | 45 weeks off ART | $5.2 \times 10^6$ |
| RM29 | SIVmac239-nef-stop/IV/200 | 73 weeks off ART | $2.0 \times 10^6$ |
| RM26<br>RM37<br>RM38<br>RM39<br>RM40 | SIVmac239-nef-stop/IV/200 | Pre-infection, weeks 1, 2, and 5 of acute infection, week 13 on ART, and weeks 3, 12, and 33 post-ART interruption | Figure 2E |
| RM2 | SIVmac251/IV/300 | Pre-infection, Day 7, 15, 32, and week 6, 12, 26, and 31 post-inoculation | Figure 2F |
| RM41 | SIVmac251/IV/300 | Pre-infection, Day 7, 15, 32, and week 6, 12, and 26 post-inoculation | Figure 2F |
| RM23<br>RM24 | SIVmac239/IV/1000 | Pre-infection, Day 9, 21, and 38 post-inoculation | Figure 2G |
| RM42 | SIVmac239/IV/1000 | Pre-infection, Day 10, 25, and 42 post-inoculation | Figure 2G |

$TCID_{50}$, median tissue culture infectious dose; IV, intravenous; ART, antiretroviral therapy

**Supplementary Table 6.** Rhesus macaques used for *in-vitro* ITS103.01LS-F(ab')$_2$ primary cells binding studies.

| | Macaque designation | Fresh/Cryo | Plasma viral load (copies/mL) | Weeks of untreated infection | ART status | SIV strain | Binding specificity |
|---|---|---|---|---|---|---|---|
| PBMC | RM14 | Fresh | $1 \times 10^4$ | 53 | ART naïve | SIVmac239 | 0.62 |
| | | Fresh | $3.2 \times 10^4$ | 87 | | | 0.8 |
| | RM15 | Fresh | $3 \times 10^6$ | 75 | ART naïve | SIVmac239x | 0.95 |
| | RM16 | Fresh | $1.4 \times 10^8$ | 10 | ART naïve | SIVmac239 | 0.77 |
| | RM9 | Cryo | - | Uninfected control | - | - | 0.83 |
| | RM18 | Fresh | - | Uninfected control | - | - | 0.88 |
| | RM34 | Fresh | - | Uninfected control | - | - | 1.02 |
| Axillary LNMC | RM16 | Fresh | $1.4 \times 10^8$ | 10 | ART naïve | SIVmac239 | 1.08 |
| | RM43 | Cryo | $5.9 \times 10^2$ | 34 | off ART | SIVmac251 | 0.7 |
| Inguinal LNMC | RM15 | Cryo | $3.9 \times 10^7$ | 105 | ART naïve | SIVmac239x | 1.07 |
| | RM25 | Cryo | $2.6 \times 10^6$ | 12 | off ART | SIVmac239-nef-stop | 1.58 |
| | RM28 | Cryo | $2.6 \times 10^3$ | 12 | off ART | SIVmac239-nef-stop | 1.9 |
| | RM44 | Cryo | $5.5 \times 10^4$ | 12 | off ART | SIVmac239-nef-stop | 1.21 |
| Mesenteric LNMC | RM10 | Cryo | $2.7 \times 10^8$ | 12 | ART naïve | SIVmac239 | 0.76 |
| | RM16 | Fresh | $1.4 \times 10^8$ | 10 | ART naïve | SIVmac239 | 0.91 |
| | | Cryo | | | | | 1.39 |
| Spleen | RM10 | Cryo | $2.7 \times 10^8$ | 12 | ART naïve | SIVmac239 | 0.75 |
| | RM15 | Cryo | $3.9 \times 10^7$ | 105 | ART naïve | SIVmac239x | 1.77 |
| | RM16 | Fresh | $1.4 \times 10^8$ | 10 | ART naïve | SIVmac239 | 0.57 |
| | | Cryo | | | | | 0.66 |
| | RM27 | Cryo | $2.6 \times 10^6$ | 3.4 | off ART | SIVmac239-nef-stop | 0.96 |
| | RM28 | Cryo | $6.2 \times 10^5$ | 90 | off ART | SIVmac239-nef-stop | 0.77 |

PBMC, peripheral blood mononuclear cells; LNMC, lymph node mononuclear cells; ART, antiretroviral therapy

**Supplementary Table 7.** SIV-infected rhesus macaques used for ITS103.01LS-F(ab')$_2$ autoradiography studies.

| Tissue | Macaque designation | Plasma viral load (copies/mL) | Weeks of untreated infection | ART status | SIV strain | Binding specificity |
|---|---|---|---|---|---|---|
| Axillary lymph node | RM11 | 1.1 x $10^7$ | 32 | ART naïve | SIVmac239 | 1.2 |
| | RM15 | 3.9 x $10^7$ | 105 | ART naïve | SIVmac239x | 1.1 |
| | RM16 | 1.4 x $10^8$ | 10 | ART naïve | SIVmac239 | 1 |
| | RM25 | 5.2 x $10^6$ | 45 | off ART | SIVmac239-nef-stop | 0.9 |
| | RM12 | - | Uninfected control | - | - | 0.8 |
| Colon | RM4 | 1.2 x $10^4$ | 254 | ART naïve | SIVmac239 | 0.3 |
| | RM13 | - | Uninfected control | - | - | 0.8 |
| Inguinal lymph node | RM2 | 2.1 x $10^5$ | 87 | ART naïve | SIVmac251 | 0.7 |
| | RM10 | 4.7 x $10^7$ | 6 | ART naïve | SIVmac239 | 0.9 |
| | RM14 | 1 x $10^4$ | 53 | ART naïve | SIVmac239 | 0.8 |
| | RM15 | 4.7 x $10^6$ | 75 | ART naïve | SIVmac239x | 0.6 |
| | RM16 | 1.4 x $10^7$ | 6 | ART naïve | SIVmac239 | 1.7 |
| | | 1.4 x $10^8$ | 10 | | | 0.9 |
| | RM12 | - | Uninfected control | - | - | 1.1 |
| | RM13 | - | Uninfected control | - | - | 1.3 |
| | RM18 | - | Uninfected control | - | - | 0.7 |
| Jejunum | RM4 | 1.2 x $10^4$ | 254 | ART naïve | SIVmac239 | 1.2 |
| | RM13 | - | Uninfected control | - | - | 1.1 |
| Ileum | RM4 | 1.2 x $10^4$ | 254 | ART naïve | SIVmac239 | 0.9 |
| | RM13 | - | Uninfected control | - | - | 0.9 |
| Spleen | RM10 | 2.7 x $10^8$ | 12 | ART naïve | SIVmac239 | 0.8 |
| | RM15 | 3.9 x $10^7$ | 105 | ART naïve | SIVmac239x | 0.7 |
| | RM16 | 1.4 x $10^8$ | 10 | ART naïve | SIVmac239 | 0.9 |
| | RM2 | 2.1 x $10^5$ | 87 | ART naïve | SIVmac251 | 1.1 |
| | RM25 | 5.2 x $10^6$ | 45 | off ART | SIVmac239-nef-stop | 1.1 |
| | RM13 | - | Uninfected control | - | - | 0.8 |
| | RM45 | - | Uninfected control | - | - | 0.9 |

ART, antiretroviral therapy
